# c-Jun Overexpressing CAR-T Cells are Exhaustion-Resistant and Mediate Enhanced Antitumor Activity

**DOI:** 10.1101/653725

**Authors:** Rachel C. Lynn, Evan W. Weber, David Gennert, Elena Sotillo, Peng Xu, Zinaida Good, Hima Anbunathan, Robert Jones, Victor Tieu, Jeffrey Granja, Charles DeBourcy, Robbie Majzner, Ansuman T. Satpathy, Stephen R. Quake, Howard Chang, Crystal L. Mackall

## Abstract

CAR T cells mediate antitumor effects in a small subset of cancer patients, but dysfunction due to T cell exhaustion is an important barrier to progress. To investigate the biology of exhaustion in human T cells expressing CAR receptors, we used a model system employing a tonically signaling CAR, which induces hallmarks of exhaustion described in other settings. Exhaustion was associated with a profound defect in IL-2 production alongside increased chromatin accessibility of AP-1 transcription factor motifs, and overexpression of numerous bZIP and IRF transcription factors that have been implicated in inhibitory activity. Here we demonstrate that engineering CAR T cells to overexpress c-Jun, a canonical AP-1 factor, enhanced expansion potential, increased functional capacity, diminished terminal differentiation and improved antitumor potency in numerous *in vivo* tumor models. We conclude that a functional deficiency in c-Jun mediates dysfunction in exhausted human T cells and that engineering CAR T cells to overexpress c-Jun renders them exhaustion-resistant, thereby addressing a major barrier to progress for this emerging class of therapeutics.

---

Chimeric antigen receptor (CAR) T cells demonstrate impressive response rates in B cell malignancies, but long-term disease control occurs in only approximately 50% of patients with B-ALL^1^ and large B cell lymphoma^2^, and is even less frequent in CLL^3^. Moreover, despite numerous trials, CAR T cells have not mediated sustained antitumor effects in solid tumors^4^. Numerous factors limit the efficacy of CAR T cells, including heterogeneous antigen expression and a requirement for high antigen density for optimal CAR function enabling rapid selection of antigen loss variants^5–7^, the suppressive tumor microenvironment^8^ and intrinsic T cell dysfunction as a result of T cell exhaustion^3, 9,10^. T cell exhaustion has been increasingly incriminated as a cause of T cell dysfunction in CAR T cells. Tonic antigen-independent signaling, due to scFv aggregation, commonly occurs in T cells expressing CARs and can induce rapid exhaustion^9^. Integration of the CD28 endodomain into second generation CAR T cell receptors enhances expansion, but also predisposes CAR T cells to exhaustion, both in the setting of tonically signaling receptors and in CD19-28z CAR T cells exposed to high tumor burdens^9^. Increased frequency of T cells bearing exhaustion characteristics contained within CD19-BBz CAR grafts were recently demonstrated to distinguish non-responding from responding patients treated for CLL^3^. Recent studies have also suggested that engineering CAR T cells to be less susceptible to exhaustion can enhance antitumor effects, as reported following integration of the CD19-28z CAR into the TRAC locus, which apparently tuned the activating signal and thus prevented exhaustion^10^, and as suggested in a recent case report, whereby apparent cure via one clonal CD19-CAR T cell was associated with insertional mutagenesis mediated disruption of TET2^11^. Together, a broad base of data from diverse studies implicates intrinsic T cell dysfunction due to T cell exhaustion as a major factor limiting the efficacy of CAR T cell therapeutics and raises the prospect that engineering exhaustion-resistant CAR T cells could substantially improve clinical outcomes.

T cell exhaustion is characterized by overexpression of cell surface receptors that mediate inhibitory signals and widespread transcriptional and epigenetic alterations^12–16^, although a comprehensive understanding of the mechanisms responsible for impaired function in exhausted T cells is lacking. PD-1 blockade can reinvigorate some exhausted T cells^17^, but is unable to fully restore function, and trials using PD-1 blockade to prevent or reverse exhaustion in CAR T cells have not yet demonstrated efficacy^18^. Using a model wherein healthy T cells are driven to exhaustion via expression of a tonically signaling CAR, we observed that exhausted human T cells rapidly developed a profound defect in IL-2 production. This phenotype was associated with widespread epigenomic dysregulation of AP-1 transcription factor binding motifs and overexpression of numerous bZIP and IRF family transcription factors that have been implicated in inhibitory AP-1 signaling. Therefore, we tested the hypothesis that dysfunction in this setting resulted from an imbalance between activating and inhibitory bZIP/IRF transcriptional activity, which could be rectified by enforced expression of c-Jun, an AP-1 family transcription factor associated with productive T cell activation. Consistent with this hypothesis, c-Jun overexpression rendered CAR T cells exhaustion-resistant, as demonstrated by enhanced expansion potential *in vitro* and *in vivo*, increased functional capacity, diminished terminal differentiation and improved antitumor potency in numerous models.

## RESULTS

### Expression of the HA-28z CAR in human T cells rapidly induces hallmark features of T cell exhaustion

We previously described development of an exhausted phenotype in human T cells following expression of a CAR incorporating the GD2-specific 14g2a scFv, TCR zeta and a CD28 endodomain (GD2-28z CAR), as a result of tonic signaling mediated via antigen-independent aggregation^9^. Here we demonstrate that expression of a CAR containing the 14g2a scFv bearing the E101K point mutation, which renders a higher affinity (HA) interaction with GD2^19^ (HA-28z CAR), similarly induces exhaustion in human T cells albeit with a more severe phenotype (Fig. 1 and Extended Data Fig. 1a-c). In contrast to CD19-28z CAR T cells, HA-28z CAR T cells displayed profound phenotypic and functional hallmarks of exhaustion, including reduced expansion in culture (Fig. 1a), increased surface expression of the inhibitory receptors PD-1, TIM-3, LAG-3, and CD39 (Fig. 1b, Extended Data Fig. 1d), exaggerated effector differentiation and poor memory formation (Fig. 1c, Extended Data Fig. 1e), and diminished IFN-γ and markedly decreased IL-2 production when stimulated with CD19^+^GD2^+^ Nalm6 leukemia (Fig. 1d).

**Figure 1:**
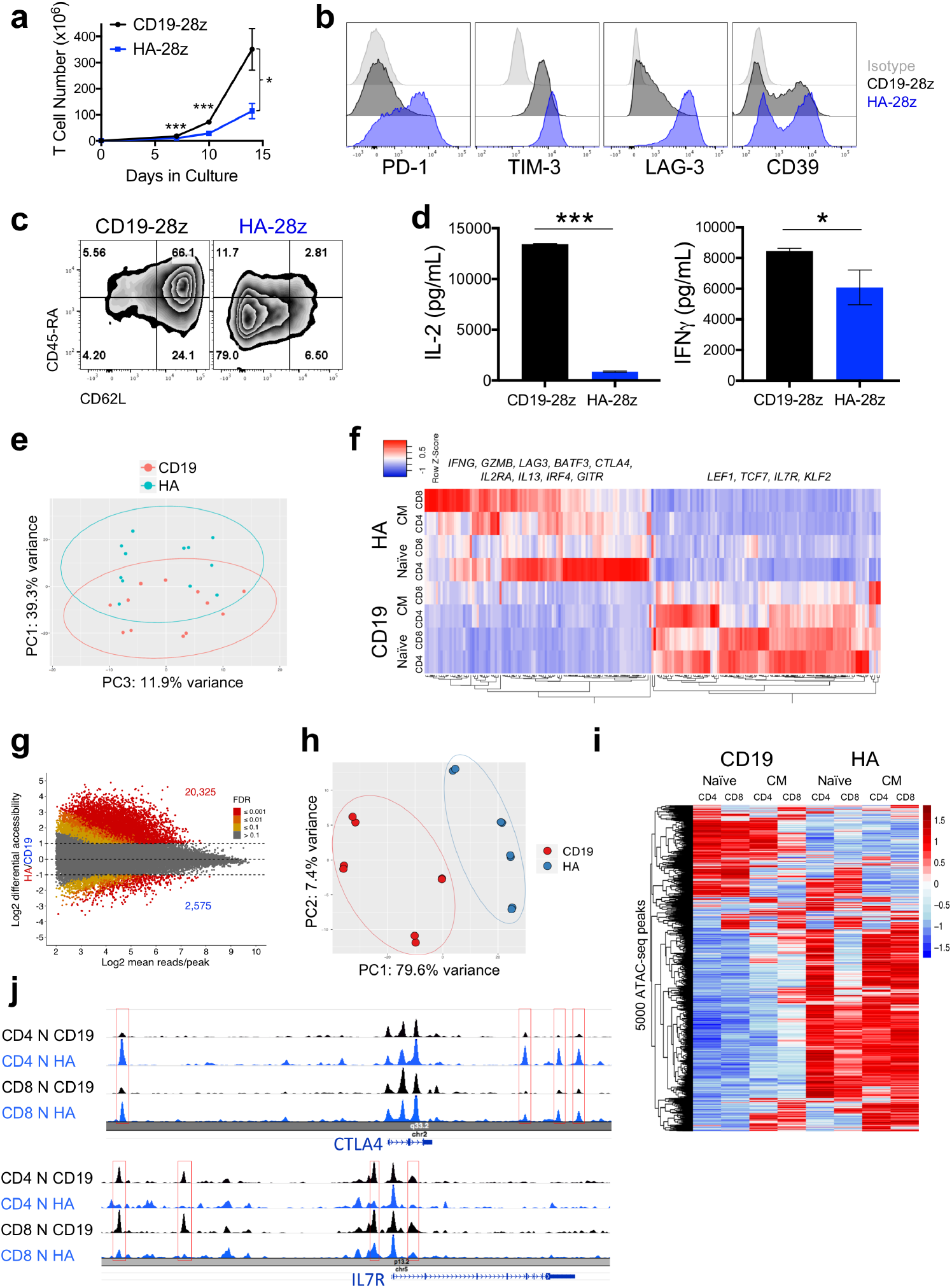
HA-28z CAR T cells manifest phenotypic, functional, transcriptional and epigenetic hallmarks of T cell exhaustion. **a)** Decreased expansion of HA-28z vs CD19-28z CAR T cells during primary expansion culture. D0=bead activation, D2=transduction. Error bars represent mean ± SEM from n=10 donors. **b)** Surface expression of exhaustion associated markers (D10). **c)** CD19-28z primarily comprise T stem cell memory (CD45RA^+^CD62L^+^) and central memory (CD45RA^-^ CD62L^+^, whereas HA-28z primarily comprise CD45RA^−^CD62L^−^ effector memory cells (D10). **d)** IL-2 (left) and IFNγ (right) release following 24-hour co-culture with CD19^+^GD2^+^ Nalm6-GD2 leukemia cells. Error bars represent mean ± SD from triplicate wells. One representative donor shown for each assay. **e)** Principle component analysis (PCA) of global transcriptional profiles of Naïve- and CM-derived CD19 or HA CAR T cells at days 7, 10, and 14 in culture. PC1 (39.3% variance) separates CD19 from HA CAR T cells. **f)** Gene expression of the top 200 genes driving PC1. Genes of interest in each cluster are listed above. **g)** Differentially accessible chromatin regions in CD8^+^ CD19 and HA-28z CAR T cells (D10). Both N and CM subsets are incorporated for each CAR. **h)** PCA of ATAC-seq chromatin accessibility in CD19 or HA-28z CAR T cells (D10). PC1 (76.9% variance) separates CD19 from HA CAR samples. **i)** Global chromatin accessibility profile of subset-derived CD19 and HA-28z CAR T cells (D10). Top 5000 differentially accessible regions (peaks). **j)** Differentially accessible enhancer regions in CD19 and HA CAR T cells in the *CTLA4* (top) or *IL7R* (bottom) loci. N – naïve, CM – central memory. * p < .05, ** p < .01, *** p < .001. ns p >.05.

To better elucidate the molecular underpinnings of T cell exhaustion in this system, we interrogated the transcriptome of HA-28z compared to CD19-28z CAR T cells. We transduced purified naïve (N) and central memory (CM) T cells with HA or CD19-28z CAR then isolated RNA on days 7, 10, and 14 of culture. Sorting pre-selected subsets allowed us to assess the impact of T cell differentiation state and the distinction between CD4 and CD8 exhaustion in the development of T cell exhaustion in this model. Principle component analysis (PCA) across all 24 samples revealed that the strongest driver of variance was the presence of the HA- vs CD19-28z CAR (PC1, 39.3% variance, Fig. 1e), consistent with a model wherein tonic signaling in HA-28z CAR T cells drives exhaustion in all T cell subsets studied. Distinctions were, however, observed based upon the starting differentiation state, since N vs CM was reflected in PC2 (22.88% variance) (Extended Data Fig1f) and between CD4 vs CD8 populations, which drove PC3 (11.9% variance) (Fig. 1e and Extended Data Fig. 1f).

Among the top 200 genes driving PC1 (most differentially expressed in HA- vs CD19-28z CAR T cells across all subsets) (Fig. 1f) were genes associated with activation (*IFNG*, *GZMB*, *IL2RA*), inhibitory receptors (*LAG3*, *CTLA4*) and some inflammatory chemokines/cytokines (*CXCL8*, *IL13*, *IL1A*), whereas genes downregulated in HA-28z CAR T cells include genes associated with naïve and memory T cells (*IL7R*, *TCF7*, *LEF1*, and *KLF2*). Using GSEA we demonstrated that genes upregulated in day 10 HA-28z vs CD19-28z CAR T cells overlap with exhaustion-associated gene sets previously described in chronic LCMV mouse models^13^ (Extended Data Fig. 1g). Although the degree of exhaustion in GD2-28z CAR T cells is less profound, differential gene expression analysis of single cell GD2-28z vs CD19-28z CAR T cells revealed a similar gene expression profile (Extended Data Fig. 2). Together, these data credential HA-28z and GD2-28z expressing T cells as models for the study of human T cell exhaustion.

T cell exhaustion is associated with changes in chromatin accessibility in mouse models and human patients with chronic viral infections and cancer^12, 17^. Chromatin accessibility analyses using ATAC-seq^20^ (Extended Data Fig. 3) of N or CM derived CD4^+^ and CD8^+^ HA-28z vs CD19-28z CAR T cells demonstrated significant changes in the epigenetic signature on day 10 of culture (Fig. 1g) with CD8^+^ HA-28z CAR T cells displaying >20,000 unique chromatin accessible regions (peaks) compared to < 3,000 unique peaks in CD8^+^ CD19-28z CAR T cells (FDR < 0.1 and log2FC > 1). These patterns of changes in exhaustion-induced chromatin accessibility were similar in CD4^+^ T cells (Extended Data Fig. 4a). Similar to the transcriptomic analysis, PCA revealed HA- vs. CD19-CAR as the strongest driver of differential chromatin states (PC1 variance 79.6%, Fig. 1h), with weaker but significant differences observed between N vs CM cells (PC2 variance 7.4%), and between the CD4 vs CD8 subsets (PC3 variance 6.5%)(Extended Data Fig. 4b). Clustering the top 5000 differentially accessible regions (peaks) revealed globally similar chromatin accessibility in HA-28z CAR T cells regardless of starting subset (Fig. 1i). As expected, HA-28z CAR T cells demonstrated increased chromatin accessibility in regulatory sites near exhaustion-associated genes such as *CTLA4*, and decreased accessibility in regulatory sites near memory associated genes such as *IL7R* (Fig. 1j).

### Epigenetic and transcriptional analyses reveal a strong AP-1 signature in exhausted CAR T cells

To identify transcriptional programs predicted to be dysregulated by the epigenetic changes induced in exhausted T cells, we compared transcription factor (TF) motif deviation in exhausted vs healthy CAR T cell open chromatin. Using ChromVAR analysis^21^, we identified the 25 most differential motifs across all 8 samples and found that many of these belong to the AP-1(bZIP) family (Fig. 2a). Similarly, TF motif enrichment analysis revealed AP-1/bZIP and bZIP/IRF binding motifs as among the most significantly enriched in exhausted CAR T cells (Fig. 2b and Extended Data Fig. 4c).

**Figure 2:**
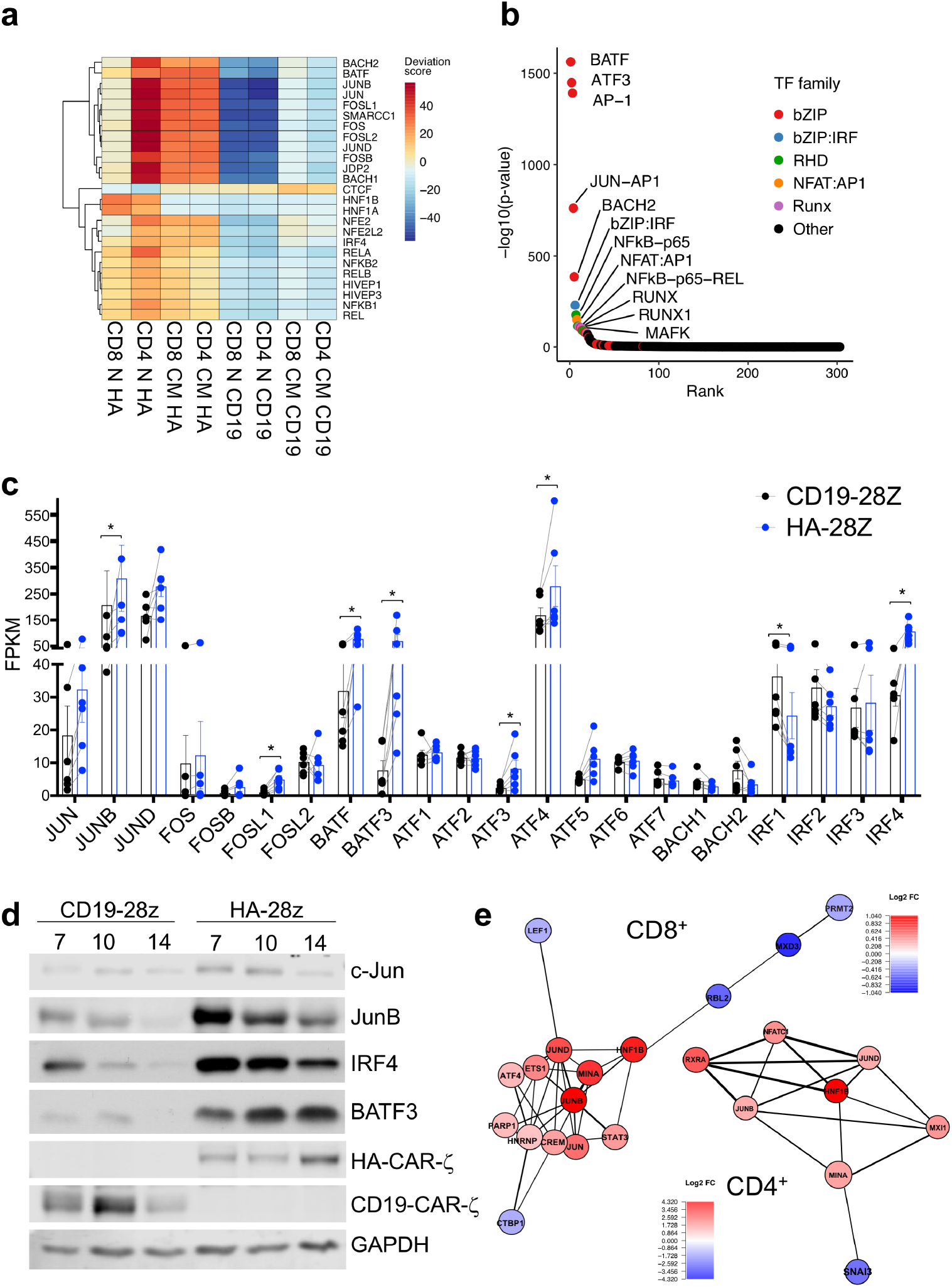
AP-1 family signature in exhausted CAR T cells. **a)** Top 25 transcription factor motif deviation scores in Day 10 HA vs CD19-CAR expressing T cells by chromVAR analysis reveal numerous AP-1(bZIP) family members in CD4^+^ and CD8^+^ T cells derived from N or CM subsets (D10). **b)** TF motif enrichment analysis in N CD8^+^ HA-28z CAR T cells demonstrates AP-1 (bZIP) family motifs as the most significantly enriched. **c)** Bulk RNA-seq expression (FPKM) of indicated AP-1 (bZIP) and IRF family members in CD19 (black) and HA-28z (blue) CAR T cells. Error bars represent mean ± SEM from n=6 samples across 3 donors showing paired CD19 vs HA expression for each gene. p-values were generated using the Wilcoxon matched-pairs signed rank test. **d)** CD19-28z and HA-28z CAR T cells were lysed and expression of the indicated AP-1 family proteins was assessed by western blot. Increased protein expression of JunB, BATF3, and IRF4 in HA-28z CAR T cells compared to CD19 CAR T cells was confirmed at days 7, 10, and 14 of culture. **e)** Correlation network of exhaustion-related transcription factors in N-derived CD8^+^ (left) and CD4^+^ (right) GD2-28z CAR T cells using single cell RNA-seq analysis. Transcription factor genes identified as differentially expressed (p < 0.05) by DESeq2 form the nodes of the network. Colors represent log2 fold-change (FC) (GD2 vs CD19 CAR). Edge thickness represents the magnitude of correlation in expression between the relevant pair of genes across cells. A correlation score > 0.1 was used to construct networks. * p < .05, ** p < .01, *** p < .001. ns p >.05.

Clustering differentially accessible peaks by shared TF motif enrichment identified 4 clusters associated with exhausted HA-28z CAR T cells (Extended Data Fig. 4d, EX1-EX4). Exhaustion-associated clusters contained peaks in the vicinity of genes like *BTLA*, *CD39*, *IFNG*, and *CTLA4,* suggesting common TF regulation of exhaustion-associated genes. All 4 exhaustion-associated clusters displayed strong enrichment for AP-1 and AP-1-related family TFs, implicating widespread AP-1 TF modulation of exhaustion-associated gene regulation. We also observed strong enrichment for NFκB, NFAT, and RUNX TF family motifs in some of the exhaustion clusters, indicating that a subset of exhaustion-related genes may be regulated by these transcriptional programs and reproduce epigenetic signatures of exhaustion observed in other models^12, 17, 22^. Interestingly, the clusters associated with healthy CD19-28z CAR T cells (HLT1-2) showed a similar profile to a cluster strongly associated with Naïve starting subset. This observation is consistent with the idea that healthy CAR T cells maintain an epigenetic signature more closely resembling naïve-derived T cells, a subset associated with increased persistence and efficacy in adoptive T cell therapy^23^, whereas chronic antigen stimulation results in broad divergent epigenetic reprogramming.

AP-1 related TFs coordinate to form a diverse set of homo and heterodimers through interactions in the common bZIP domain and can dimerize with IRF transcription factors^14, 24^. AP-1 factor complexes compete for binding at DNA elements containing core TGA-G/C-TCA consensus motifs. Activating complexes such as those comprising the classically described AP-1 heterodimer c-Fos and c-Jun drive IL-2 transcription. Conversely, other AP-1 and IRF family members can directly antagonize c-Jun activity and/or drive immunoregulatory gene expression in T cells^14, 24–29^. To assess whether the changes in AP-1 binding chromatin accessibility were associated with increased availability of activating and inhibitory bZIP and IRF TFs, we compared transcript levels of bZIP and IRF family members using RNA-seq in exhausted HA-28z vs. healthy CD19-28z CAR T cells. Paired RNA-seq analysis across 3 different donors revealed a consistent pattern of bZIP and IRF family member overexpression, which was most significant for *JUNB*, *FOSL1, BATF, BATF3*, *ATF3, ATF4* and *IRF4* (Fig. 2c and Extended Data Fig 5a). Western blot analysis confirmed sustained protein overexpression of JunB, IRF4, and BATF3 in HA- vs. CD19 CAR T cells (Fig. 2d and Extended Data Fig 5b), with the immunoregulatory BATF/IRF TFs showing higher levels of expression compared to c-Jun. The biological significance of increased levels of inhibitory bZIP/IRF family members was further suggested by the demonstration that western blotting of Jun immunoprecipitates (IP) revealed that several inhibitory family members are in direct complex with c-Jun and JunB in HA-28z exhausted CAR T cells (Extended Data Fig. 5c). Comparison of TF profiles by single cell RNA-seq analysis of CD8^+^ T cells expressing CD19-28z vs. GD2-28z CAR confirmed that the bZIP family members *JUN*, *JUNB*, *JUND*, and *ATF4* were among the most differentially expressed and broadly connected in exhausted GD2-28z CAR T cell networks (Fig. 2e and Extended Data Fig. 2).

### c-Jun overexpression (OE) reduces functional exhaustion in CAR T cells

Based upon the evidence that exhausted CAR T cells manifest very poor IL-2 production (Fig. 1d) and preferentially overexpress bZIP and IRF transcription factors that drive immunoregulatory and exhaustion-associated programs, we hypothesized that T cell dysfunction in exhausted cells might be due to a *relative* deficiency in c-Jun/c-Fos heterodimers necessary to drive IL-2 transcription. To test this, we co-transduced HA-28z and CD19-28z CAR T cells with a bicistronic lentiviral vector to overexpress c-Jun and c-Fos. HA-28z CAR T cells engineered to overexpress AP-1 demonstrated increased IL-2 production upon antigen stimulation (Extended Data Fig. 6a-c). However, using single expression vectors we only observed enhanced functionality upon c-Jun OE in HA-28z CAR T cells, whereas transduction with a c-Fos single expression vector yielded no consistent functional improvement (Extended Data Fig. 6d-e).

To further investigate the potential for c-Jun to enhance the function of exhausted CAR T cells and to ensure constitutive c-Jun expression in all CAR expressing T cells, we created bicistronic vectors to co-express c-Jun and CAR transgenes separated by the viral P2A skipping peptide (JUN-CARs, Fig. 3a). These expression vectors increased c-Jun expression in both CD19 and HA-28z CAR T cells (Fig. 3b), although c-Jun was preferentially activated (phosphorylated) in JUN-HA CAR T cells (Fig. 3c), consistent with c-Jun N-terminal phosphorylation (JNP) by JNK proteins activated downstream of the tonic TCR signaling cascade propagated via the HA-28z CAR^30^. Upon stimulation with GD2^+^ tumor cell lines, JUN-HA-28z CAR T cells demonstrated a remarkable increase in IL-2 and IFNγ production compared to control HA-28z CAR T cells (Fig. 3d-e). The fold increase in cytokine production in the setting of c-Jun OE was substantially greater for JUN-HA CAR compared to JUN-CD19 CAR T cells (Extended Data Fig 7a-b). Similarly, JUN-HA CAR T cells demonstrated increased frequencies of SCM/CM vs E/EM subsets compared to HA-28z CAR T cells (Fig. 3f), whereas no significant difference in subset composition was observed between CD19 and JUN-CD19 CAR T cells at day 10 of culture. Together, the data are consistent with a model wherein c-Jun OE is functionally more significant in exhausted T cells, which overexpress inhibitory bZIP and IRF TFs.

**Figure 3:**
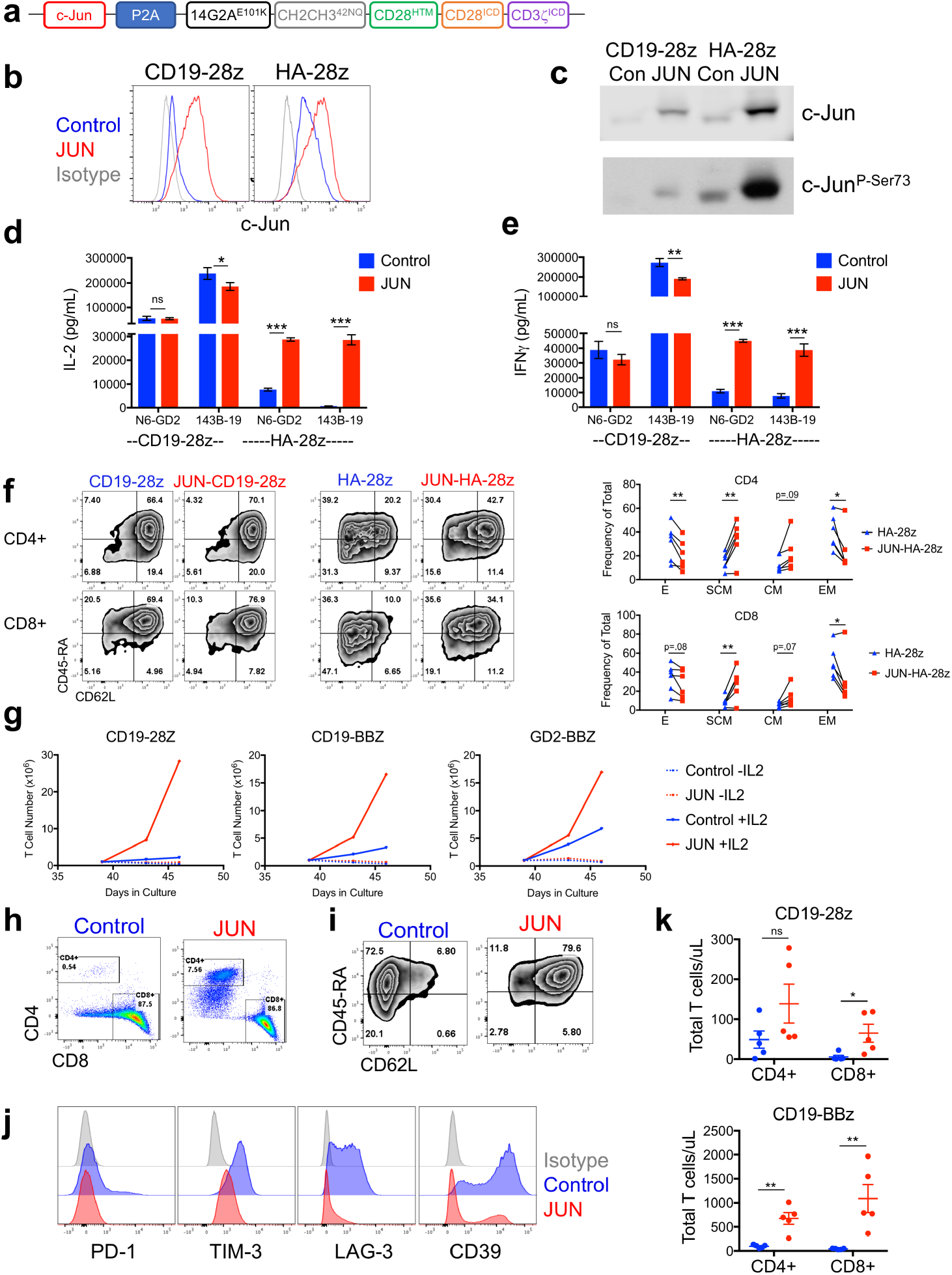
c-Jun overexpression enhances the function of exhausted CAR T cells. **a)** Schematic of the JUN-P2A-CAR expression vector. **b)** Intracellular flow cytometry demonstrating total c-Jun expression in control (blue) and JUN-modified (red) CAR T cells (D10). **c)** Western blot for total c-Jun and phospho-c-Jun^Ser73^ in control and JUN-modified CD19 and HA CAR T cells (D10). **(d)** IL-2 and **(e)** IFNγ production following 24hr co-culture of control (blue) or JUN-modified (red) CD19 and HA CAR T cells in response to antigen+ tumor cells. Error bars represent mean ± SD of triplicate wells. Data from one representative donor shown. Fold increases in IL-2 or IFNγ production in JUN vs control CAR T cells across multiple donors can be found in Extended Data Figure 7. **f)** Left: Flow cytometry plots showing representative CD45RA/CD62L expression in Control vs JUN-CAR T cells (D10). Right: Relative frequency of effector (E, RA+62L-), stem cell memory (SCM, RA+62L+), central memory (RA-62L+), and effector memory (RA-62L-) in CD4^+^ (upper) or CD8^+^ (lower) Control or JUN-HA-28z CAR T cells. n=6 donors from independent experiments. Lines indicate paired samples from the same donor. Paired, two-tailed t-tests were performed. **g)** On day 39 of culture, 1×10^6^ viable T cells from Ext Data Fig 7c were re-plated and cultured for 7 days with or without IL-2. **h-j)** Cell surface phenotype of control or JUN-CD19-28z CAR T cells from (g) on day 46. **h)** CD4 vs CD8 expression. **i)** Late expanding CD8^+^ JUN-modified CD19-28z CAR T cells have a stem cell memory phenotype (CD45RA^+^CD62L^+^). **j)** Late expanding CD8^+^ JUN-modified CD19-28z CAR T cells have reduced exhaustion marker expression compared to controls. **k)** T cells from g were cryopreserved on D10, thawed and rested overnight in IL-2. Healthy NSG mice were infused with 5×10^6^ control or JUN-modified CD19-28z or CD19-BBz CAR T cells via IV injection. On day 25 post infusion, peripheral blood T cell numbers were quantified by flow cytometry. Error bars represent mean ± SEM of n=5 mice per group. * p < .05, ** p < .01, *** p < .001. ns p >.05. HTM – hinge/transmembrane. ICD – intracellular domain.

To assess whether c-Jun OE enhances long-term proliferative capacity, which is associated with antitumor effects in solid tumors^31^, and to test whether c-Jun OE could augment function in CAR T cells without tonic signaling (CD19-28z, CD19-BBz) or in those with lesser levels of tonic signaling (GD2-BBz), we measured *in vitro* expansion of JUN-CAR T cells from 3 different healthy donors over a protracted period (Extended Data Fig. 7c). We observed a consistent pattern of enhanced long-term proliferative capacity in the presence of c-Jun OE, which remained IL-2-dependent, as these cells immediately ceased expansion in the absence of IL-2 (Fig. 3g). Consistent with c-Jun’s capacity to induce resistance to exhaustion, late expanding CD8^+^ JUN-CD19-28z CAR T cells displayed diminished expression of exhaustion markers and maintained a robust subset of cells bearing a stem cell memory (SCM) phenotype (CD45RA^+^CD62L^+^) compared to control CD19-28z CAR T cells tested at the same timepoint (Fig. 3h-j). We also evaluated homeostatic expansion of JUN-CAR T cells adoptively transferred into tumor-free NSG mice. Peripheral blood T cell numbers were increased in both JUN-CD19-28z and JUN-CD19-BBz CAR T cell treated mice compared to controls 25 days post infusion (Fig. 3k), which led to accelerated GVHD in the JUN-CD19-BBz CAR T cell-treated mice. Together the data demonstrate that c-Jun OE mitigates T cell exhaustion in numerous CARs tested, including those incorporating CD28 or 4-1BB costimulatory domains, and regardless of whether exhaustion is driven by enforced long-term expansion or tonic signaling.

### Molecular requirements for c-Jun-mediated rescue of CAR T cell exhaustion

We postulated that c-Jun OE could rescue exhausted T cells by two distinct mechanisms that are not mutually exclusive. c-Jun OE could directly enhance AP-1 mediated gene transcription by increasing availability of Fos/Jun or Jun/Jun dimers or could work indirectly by disrupting inhibitory AP-1/IRF transcriptional complexes (AP-1-i)^14, 24^ that drive exhaustion-associated gene expression (Figure 4a). In order to better understand the mechanisms by which c-Jun OE mitigates T cell exhaustion, we first sought to temporally regulate c-Jun expression by fusing the destabilization domain (DD) derived from *E. coli* dihydrofolate reductase (DHFR)^32^ to the N-terminus of c-Jun (JUN-DD, Fig. 4b). The DD is stabilized in the presence of the cell permeable small molecule trimethoprim (TMP) and results in stable c-Jun expression, but in absence of TMP, the DD is destabilized, inducing proteasomal degradation of the entire fusion protein (Fig. 4c). JUN-DD CAR T cells rapidly increased c-Jun expression in the presence of TMP (1/2max at 6.76 hours following drug exposure), whereas JUN-DD rapidly became undetectable in the absence of TMP (t_1/2_ of 1.84 hours) (Fig 4d and Extended Data Fig. 8a-b). JUN-DD-CAR T cells mediated increases in IL-2 and IFNγ production only in the presence of TMP, confirming a critical role for c-Jun levels in modulating CAR T cell functionality (Fig. 4e). We reasoned that direct effects of c-Jun on transcription could be effected quickly, and therefore tested whether functional rescue would occur when c-Jun OE in HA-28z CAR T cells was restricted to the period of acute antigen stimulation (OFF**→**ON). In contrast, if c-Jun OE was necessary to compete with inhibitory AP-1 complexes (AP-1i) during induction of exhaustion, a more prolonged exposure during T cell expansion (ON**→**OFF) may be required (Figure 4f). Compared to HA-28z CAR T cells which never experienced c-Jun OE (OFF**→**OFF), both conditions mediated partial rescue, however, full rescue of IL-2 function required c-Jun OE during both T cell expansion and antigen stimulation (ON**→**ON) (Figure 4g). This finding is consistent with a model wherein c-Jun OE both directly enhances gene transcription during acute stimulation downstream of antigen encounter and also indirectly modulates molecular reprogramming during the development of exhaustion. Further consistent with the indirect model hypothesis, we observed reductions in both protein and mRNA expression of JUNB and BATF/BATF3 family members upon c-Jun overexpression (Extended Data Figure 8c-d) as well as reduction in the JunB/BATF complexes through IP (Extended Data Figure 8e).

**Figure 4:**
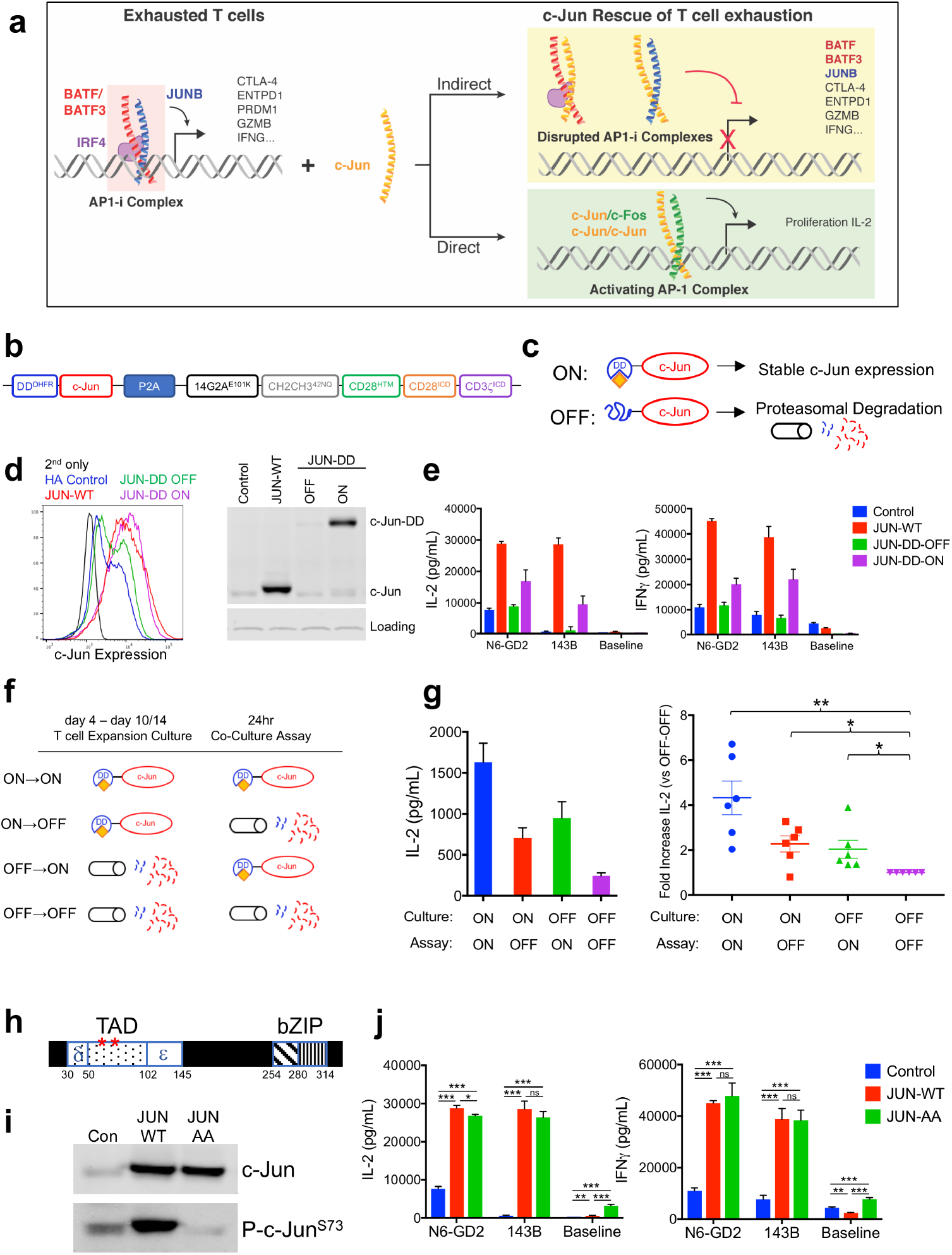
Functional rescue of exhausted HA-28z CAR T cells requires the presence of c-Jun during both chronic and acute T cell stimulation and is independent of JNP. **a)** Proposed mechanisms of c-Jun-mediated rescue of T cell exhaustion. AP-1-i indicates an inhibitory AP-1 complex. **b)** Schematic of the DD regulated JUN expression vector. **c)** Schematic of drug-induced stabilization of JUN-DD expression. Yellow diamond – TMP stabilizing molecule. **d)** Total c-Jun expression in control, JUN-WT, and JUN-DD HA-28z CAR T cells (D10) by intracellular flow cytometry (left) and western blot (right). **e)** IL-2 (left) and IFNγ (right) production in Control (blue), JUN-WT (red), or JUN-DD (OFF-green, ON-purple) modified HA-28z CAR T cells 24hr following stimulation with Nalm6-GD2 or 143B target cells, or media alone (baseline) (D10). In d-e) OFF indicates without TMP, ON indicates T cells cultured in the presence of 10uM TMP from D4 and during co-culture. In **f-g)** TMP was added either during T cell expansion (starting at D4) or only during co-culture with tumor cells as indicated in f. For ON**→**OFF and OFF**→**ON conditions, TMP was removed/added 18hr prior to co-culture to ensure complete c-Jun degradation/stabilization, respectively, prior to antigen exposure. **g)** IL-2 expression in one representative donor (left, SD across triplicate wells) and fold increase in IL-2 (SEM of n=6 independent experiments representing 3 different donors, relative to OFF-OFF condition). **h-j)** Increased functional activity of JUN-CAR T cells is independent of Jun N-terminal phosphorylation (JNP). **h)** Schematic of c-Jun protein showing N-terminal transactivation domain (TAD). Red asterisks represent the JNP sites at Ser63 and Ser73 which are mutated to alanine in the JUN-AA mutant. **i)** Western blot of total c-Jun and c-Jun-P^Ser73^ in control, JUN-WT, and JUN-AA HA-28z CAR T cells. **j)** IL-2 (left) and IFNγ (right) release in control (blue), JUN-WT (red), and JUN-AA (green) HA-28z CAR T cells following 24hr stimulation with Nalm6-GD2 or 143B target cells or media alone (Baseline). Error bars represent mean ± SD of triplicate wells. Representative of 3 independent experiments. * p < .05, ** p < .01, *** p < .001. ns p >.05. HTM – hinge/transmembrane, ICD – intracellular domain, DD – destabilizing domain from E. coli DHFR, TMP – trimethoprim, WT – wildtype.

The indirect model of c-Jun-mediated disruption of inhibitory AP-1 complexes would be independent of direct c-Jun transcriptional activation. To test whether direct c-Jun-mediated gene activation was necessary for the functional rescue of T cell exhaustion, we created a JNP-deficient c-Jun with alanine substitutions of Ser63 and Ser73 in the c-Jun transactivation domain to prevent phosphorylation at these sites (c-Jun^AA^), which has been demonstrated to be important for c-Jun mediated gene transcription^33 34^ (Figure 4h-i). Remarkably, JUN^AA^-HA-28z CAR T cells demonstrated equivalent increases in IL-2 and IFNγ production compared to wildtype JUN-CAR T cells (Figure 4j). Together, our data is consistent with a model wherein c-Jun mediated rescue of exhausted does not require direct gene activation.

### JUN-CAR T cells mediate enhanced antitumor activity *in vivo*

We next asked whether JUN-CAR T cells would demonstrate enhanced activity *in vivo*. Nalm6-GD2^+^ leukemia cells were engrafted into mice and treated with control HA-28z or JUN-HA-28z CAR T cells on day 3. While HA-28z CAR T cells exhibited some anti-tumor activity, the treatment ultimately failed as all mice succumbed to disease (median survival d59). In contrast, JUN-HA-28z CAR T cells mediated complete tumor regression by day 24 and provided long term, tumor-free survival (Fig. 5a-c). We next sought to address whether c-Jun OE could enhance the functionality of CARs targeting solid tumors. We evaluated the effect of JUN-CAR using Her2 and GD2 targeting CARs incorporating the 4-1BB costimulatory domain, which has become the preferred signaling domain for imparting long-term persistence^35–37^. In a protracted ex vivo killing assay of 143b osteosarcoma JUN-Her2-BBz CAR T cells manifested significantly more potent killing activity at a 1:8 effector:target (E:T) ratio, consistent with enhanced potency on a per cell basis (Fig. 5d-e). Similarly, JUN-Her2-BBz CAR T cells prevented tumor growth *in vivo* and led to dramatically improved long-term survival, which was associated with increased T cell expansion *in vivo* (Fig. 5f-h). Similar results were observed when comparing GD2-BBz and JUN-GD2-BBz CAR T cells against 143b osteosarcoma (Extended Data Fig. 9), confirming the benefit of c-Jun OE in CAR T cells responding to solid tumors and in CAR T cells incorporating 4-1BB signaling domains.

**Figure 5:**
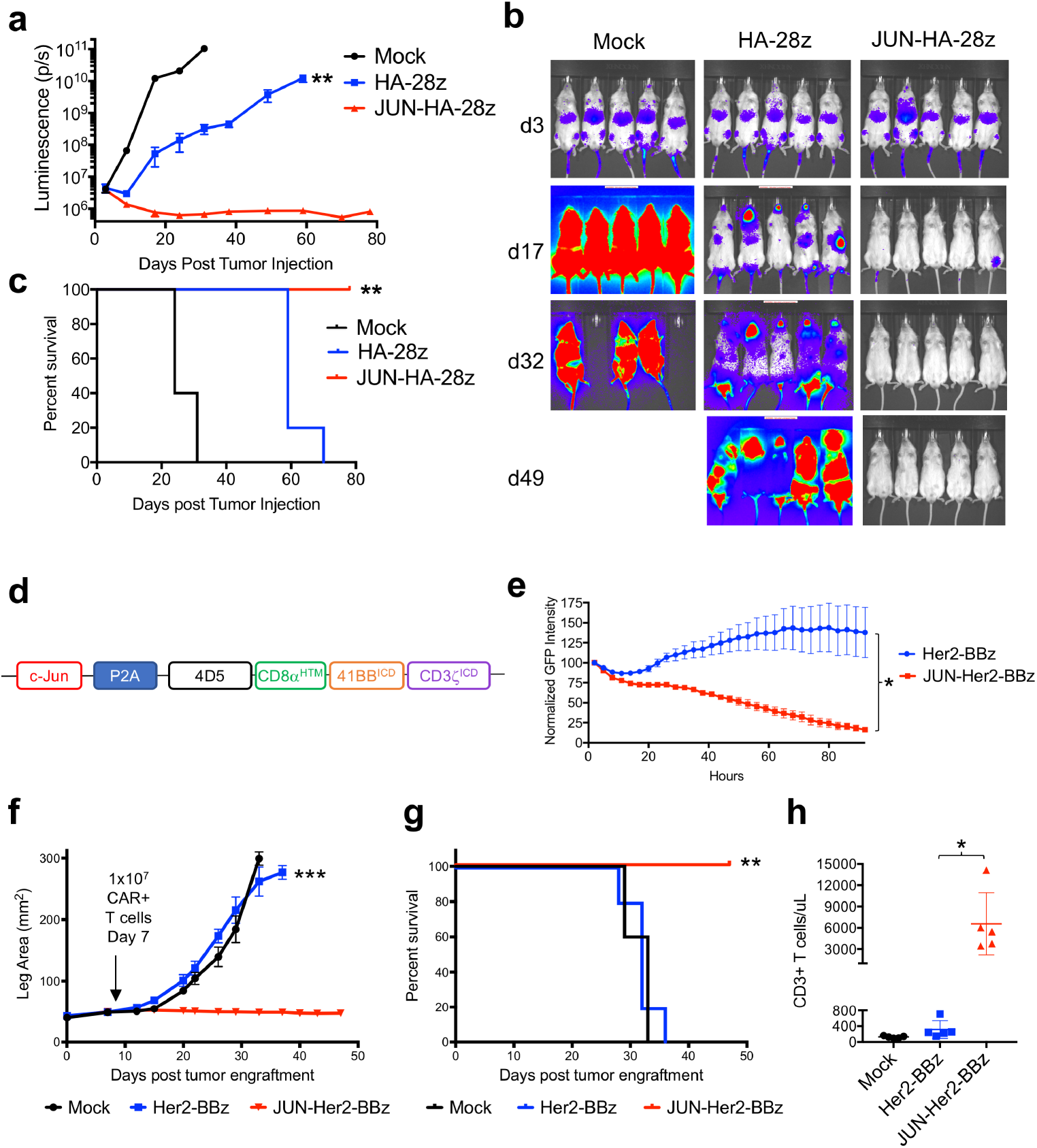
JUN-modified CAR T cells increase in vivo activity against leukemia and solid tumors. In **a-c**, NSG mice were inoculated with 1×10^6^ Nalm6-GD2 leukemia cells via IV injection. 3×10^6^ Mock, HA-28z, or JUN-HA-28z CAR+ T cells were given IV on d3. Tumor progression was monitored using bioluminescent imaging **(a-b)**. Scales are normalized for all time points. **c)** JUN-HA-28z CAR T cells induced long term tumor-free survival. Error bars represent mean ± SEM of n=5 mice/group. This finding was reproducible in >3 independent experiments, however, in some experiments long term survival was diminished due to outgrowth of GD2(-) Nalm6 clones. **d)** Schematic of JUN-Her2-BBz retroviral vector construct. **e)** Her2-BBz CAR T cell lysis of GFP+ Nalm6-Her2 target cells at 1:8 E:T ratio. Error bars represent mean ± SD of triplicate wells. Representative of 2 independent experiments. In **f-h**, NSG mice were inoculated with 1×10^6^ 143b-19 osteosarcoma cells via intramuscular injection. 1×10^7^ Mock, Her2-BBz, or JUN-Her2-BBz CAR T cells were given IV on d7. **f)** Tumor growth was monitored by caliper measurements. **g)** JUN-Her2-BBz CAR T cell treated mice maintained long term, tumor-free survival. **h)** On d20 following tumor cell implantation, peripheral blood T cells were quantified in mice treated as in f. Error bars represent mean ± SEM of n=5 mice/group. Representative of 2 independent experiments with similar results. * p < .05, ** p < .01, *** p < .001. HTM – hinge/transmembrane. ICD – intracellular domain.

### c-Jun overexpression decreases CAR T cell activation threshold and permits recognition of tumor cells with lower antigen density

The elevated levels of inhibitory AP1 family members in exhausted CAR T cells raised the prospect that dysfunction in exhausted CAR T cells relates, at least in part, to a higher threshold for activation, which might be normalized by restoring the balance of activating vs inhibitory AP1 family members. To test this, we compared cytokine production of HA-CAR vs. JUN-HA-CAR T cells in response to serial dilutions of plate-bound 1A7, an anti-idiotype antibody that binds the 14g2a scFv, allowing for control of stimulus strength. c-Jun OE substantially enhanced maximal IL-2 and IFNγ produced, and also substantially lowered the amount of 1A7 needed to induce IL-2 secretion, consistent with a reduced activation threshold in JUN-HA-CAR T cells (Fig. 6a-b).

**Figure 6:**
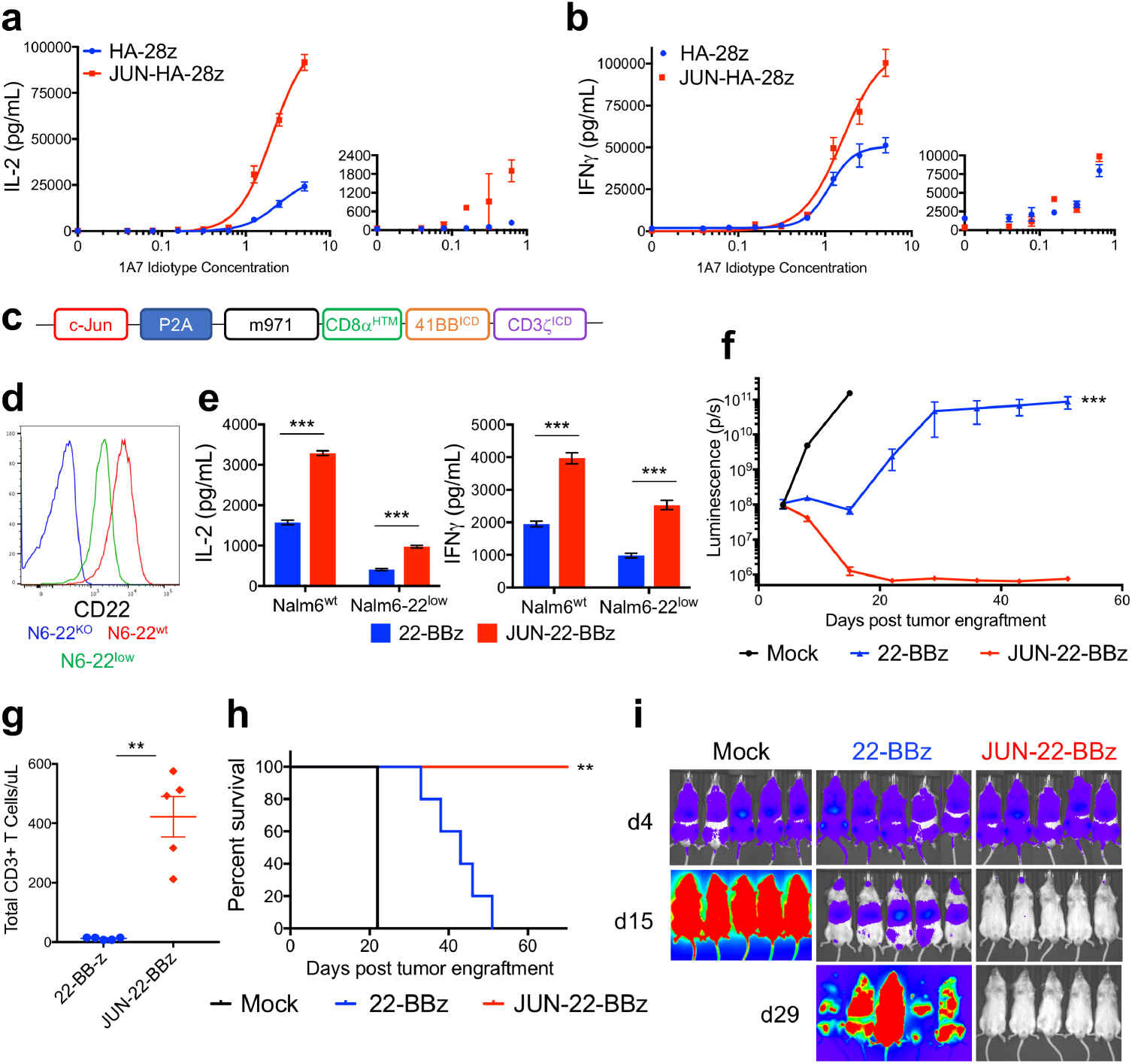
JUN-CAR T cells enhance T cell function under suboptimal stimulation. **a)** IL-2 and **b)** IFNγ production following 24hr stimulation of control (blue) or JUN-modified (red) HA-28z CAR T cells with immobilized 1A7 anti-CAR idiotype antibody. Each curve was fit with non-linear dose response kinetics to determine EC50. Smaller graphs to the right visualize curve where antibody concentration is 0-1 ug/mL. **c)** Vector schematic of JUN-CD22-BBz retroviral vector construct. **d)** CD22 surface expression on parental Nalm6 (red), Nalm6-22KO (blue), and Nalm6-22KO+CD22^low^ (green). **e)** IL-2 (left) and IFNγ (right) release following co-culture of control (blue) or JUN (red) CD22-BBz CAR T cells exposed to Nalm6 and Nalm6-22^low^. Error bars represent mean ± SD of triplicate wells. Representative of 3 independent experiments. In **f-i)**, NSG mice were inoculated with 1×10^6^ Nalm6-22^low^ leukemia cells on day 0. On day 4, 3×10^6^ control or JUN-CD22-BBz CAR^+^ T cells or 3×10^6^ Mock transduced T cells were transferred IV. Tumor growth was monitored by bioluminescent imaging **(f)** with images **(i)**. **g)** Mice receiving JUN-22BBz CAR T cells display increased peripheral blood T cells on day 23. **h)** JUN expression significantly improved long term survival of CAR treated mice. In f-g, error bars represent mean ± SEM of n=5 mice per group. Representative of 2 independent experiments with similar results. * p < .05, ** p < .01, *** p < .001. HTM – hinge/transmembrane. ICD – intracellular domain.

Limiting target antigen expression levels on tumor cells is increasingly recognized to limit CAR functionality^5, 6, 38^. We recently reported CD22^dim^ relapses in leukemia patients following initial responses to CD22 CAR therapy. Because c-Jun OE lowers the activation threshold in tonically signaling HA-28z CAR T cells, we assessed whether JUN-CARs would recognize and kill tumor cells with lower antigen density, which may escape recognition by control CAR T cells. When JUN-CD22-BBz CAR (Fig. 6c) T cells were challenged with CD22^low^ leukemia (Fig. 6d), JUN-CAR T cells exhibited increased cytokine production *in vitro* (Fig. 6e) and dramatically increased *in vivo* anti-tumor activity (Fig. 6f-i). Control CD22-BBz CAR T cells demonstrated initial activity when given at a dose of 3×10^6^ CAR T cells, but this treatment ultimately failed to control tumor growth (mean survival d45). In contrast, JUN-CD22-BBz CAR T cells mediated significant antitumor effects and were completely curative. Thus, c-Jun OE demonstrates significantly improved antitumor control in 4 tumor models, and is associated with improved expansion, resistance to exhaustion, and improved capacity to recognize low antigen targets.

## DISCUSSION

Several lines of evidence implicate exhaustion in limiting the potency of CAR T cells^3, 9–11^. Using a tonically signaling CAR capable of inducing the hallmark features of exhaustion during a controlled *in vitro* culture system, we comprehensively profiled the transcriptomic and epigenetic state of human CAR T cell exhaustion. Our findings largely align with data generated from mice and humans with cancer and chronic viral infection, implying that CAR T cell exhaustion is similar to T cell exhaustion occurring in other settings and thus validate the model system as relevant for the study of human T cell exhaustion. Our ATAC-seq data identified novel exhaustion-specific regulatory regions, which allowed us to identify transcriptional regulators, most notably of the AP-1 related bZIP/IRF families, likely responsible for driving exhaustion-associated gene expression.

Given the combined observations of a profound defect in IL-2 production, significant epigenetic dysregulation of AP-1 binding sites, and overexpression of numerous bZIP and IRF TFs implicated in inhibiting the activity of c-Jun^25–27^ and/or promoting gene expression associated with exhaustion and dysfunction^14, 24, 28, 29^, we hypothesized that diminished functionality in exhausted T cells could result from a relative deficiency in AP-1. A related hypothesis was previously put forth by Martinez et al, who suggested that exhaustion in murine T cells results from partnerless NFAT^39^. To test our hypothesis, we engineered CAR T cells to overexpress c-Jun and/or c-Fos. Remarkably, no significant effects were observed with c-Fos, whereas c-Jun OE alleviated the phenotypic and functional hallmarks of exhaustion, and substantially enhanced CAR T cell potency, not only in HA-28z tonically signaling CAR T cells, but also in clinically relevant GD2-BBz and Her2-BBz CAR T cells tested in solid tumor models. JUN-CAR T cells also demonstrated increased potency when encountering tumor cells with low antigen density, as illustrated by the enhanced capacity for JUN-CD22-BBz CAR T cells to eradicate antigen-dim leukemia.

Two distinct but non-mutually exclusive mechanisms could explain the functional rescue we observed in JUN-CAR T cells. The first postulates that overexpression of inhibitory bZIP and IRF TFs preclude c-Jun/c-Fos-mediated transcriptional activation of genes like IL-2. Thus, c-Jun overexpression leads to *direct* transcriptional activation. Using a regulatable c-Jun model, we demonstrated that full functional rescue requires the presence of c-Jun during both the T cell expansion and acute stimulation phase, suggesting c-Jun expression during chronic T cell activation alters transcriptional reprogramming during the development of exhaustion. Furthermore, the finding that the c-Jun^AA^ JNP-mutant, which should limit the transactivation potential of c-Jun, also mediated exhaustion-resistance, suggests that c-Jun overexpression may rescue function by an alternative mechanism. A second model posits that c-Jun OE acts in a dominant negative fashion by *indirectly* antagonizing IRF/BATF or other inhibitory homo- or heterodimers which mediate transcriptional activity that contributes to the exhaustion profile, as previously reported^14^. The inability for Fos overexpression to enhance function is consistent with the indirect antagonization model, as Fos has not been described to heterodimerize with BATF proteins. Further, the previously reported finding that BACH2 protects from terminal effector differentiation by blocking AP-1 sites^40^ is also consistent with an indirect model wherein c-Jun similarly acts to block access of inhibitory AP-1 complexes to enhancer regions, as terminal effector differentiation is a hallmark of exhaustion in our model. Whether through a direct or indirect mechanism, our data are consistent with the hypothesis that an imbalance of c-Jun to other bZIP/IRF TFs contributes to the biology of human T cell exhaustion, and that functional rescue can be achieved by c-Jun replacement.

The impressive effects of c-Jun overexpression in numerous preclinical tumor models raise the prospect of clinical testing of JUN-CAR T cells. c-Jun was first discovered as the cellular homolog (c-Jun) of v-Jun^41^, the viral oncogene from avian sarcoma virus and c-Jun expression has been described in cancer. However, despite extensive study, tumor promoting effects of c-Jun in human cancer are limited to *JUN* amplification in undifferentiated soft-tissue sarcomas, where it appears to prevent adipocyte differentiation^42^, and overexpression of c-Jun in Hodgkin’s Disease and anaplastic large cell lymphoma, where it may enhance proliferation and suppress apoptosis^43^. c-Jun has not been implicated as an oncogene in mature T cells, which appear to be generally resistant to transformation. Consistent with this finding, we observed that c-Jun overexpressing T cells manifest increased proliferative capacity *in vitro* and *in vivo*, but remain IL-2-dependent and we observed no evidence for leukemic transformation. *Ras*-mediated transformation in rodent models required JNP^44^, therefore, our c-Jun^AA^ mutant, which equally rescues CAR T cell function, could be implemented to mitigate theoretical oncogenic risk. Furthermore, we developed a regulatable version of c-Jun that can be induced using trimethoprim, an FDA approved small molecule. The favorable kinetics for rapid, inducible stabilization of c-Jun could allow fine-tuned control of c-Jun expression in CAR T cells in patients while mitigating long-term risk. An alternative approach to address the theoretical risks of *JUN* overexpression would be to integrate a suicide domain into JUN-CAR T cells, an approach that has already been tested in clinical trials^45^.

In summary, our findings highlight the power of a deconstructed model of human T cell exhaustion to interrogate the biology of this complex phenomena. Using this approach, we discovered a fundamental role for the AP-1/bZIP family in human T cell exhaustion and demonstrated that c-Jun OE renders CAR T cells exhaustion-resistant, which enhances their capacity to control tumor growth *in vivo*, and enhances recognition of antigen-dim targets, thus addressing major barriers to progress with this novel class of therapeutics.

## METHODS

### Viral vector construction

MSGV retroviral vectors encoding the following CARs were previously described: CD19-28z, CD19-BBz, GD2-28z, GD2-BBz, Her2-BBz, and CD22-BBz. To create the HA-28z CAR, a point mutation was introduced into the 14G2a scFv of the GD2-28z CAR plasmid to create the E101K mutation. The “4/2NQ” mutations^46^ were introduced into the CH2CH3 domains of the IgG1 spacer region to diminish Fc receptor recognition for *in vivo* use of HA-28z CAR T cells. Codon optimized cDNAs encoding c-Jun (JUN), c-Fos (FOS), and truncated NGFR (tNGFR) were synthesized by IDT and cloned into lentiviral expression vectors to create JUN-P2A-FOS, and JUN and FOS single expression vectors co-expressing tNGFR under the separate PGK promoter. JUN-P2A was then subcloned into the XhoI site of MSGV CAR vectors using the In-Fusion HD cloning kit (Takara) upstream of the CAR leader sequence to create JUN-P2A-CAR retroviral vectors. For JUN-AA, point mutations were introduced to convert Ser63 and Ser73 to Ala. The E.coli DHFR-DD sequence was inserted upstream of Jun to create JUN-DD constructs. In some cases, GFP cDNA was also subcloned upstream of the CAR to create GFP-P2A-CAR vector controls.

### Viral vector production

Retroviral supernatant was produced in the 293GP packaging cell line as previously described. Briefly, 70% confluent 293GP 20cm plates were co-transfected with 20ug MSGV vector plasmid and 10ug RD114 envelope plasmid DNA using Lipofectamine 2000. Media was replaced at 24 and 48 hours post transfection. The 48HR and 72HR viral supernatants were harvested, centrifuged to remove cell debris, and frozen at −80C for future use. Third generation, self-inactivating lentiviral supernatant was produced in the 293T packaging cell line as previously described. Briefly, 70% confluent 293T 20cm plates were co-transfected with 18ug pELNS vector plasmid, and 18ug pRSV-Rev, 18ug pMDLg/pRRE (Gag/Pol) and 7ug pMD2.G (VSVG envelope) packaging plasmid DNA using Lipofectamine 2000. Media was replaced at 24 hours post transfection. The 24HR and 48HR viral supernatants were harvested, combined, and concentrated by ultracentrifugation at 28,000 RPM for 2.5hr. Concentrated lentiviral stocks were frozen at −80C for future use.

### T cell isolation

Primary human T cells were isolated from healthy donors using the RosetteSep Human T cell Enrichment kit (Stem Cell Technologies). Buffy coats were purchased from Stanford Blood Center and processed according to the manufacturer’s protocol using Lymphoprep density gradient medium and SepMate-50 tubes. Isolated T cells were cryopreserved at 2×10^7^ T cells per vial in CryoStor CS10 cryopreservation medium (Stem Cell Technologies).

### CAR T cell production

Cryopreserved T cells were thawed and activated same day with Human T-Expander CD3/CD28 Dynabeads (Gibco) at 3:1 beads:cell ratio in T cell media (AIMV supplemented with 5% FBS, 10mM HEPES, 2mM GlutaMAX, 100 U/mL penicillin, and 100ug/mL streptomycin (Gibco)). Recombinant human IL-2 (Peprotech) was provided at 100 U/mL. T cells were transduced with retroviral vector on days 2 and 3 post activation and maintained at 0.5-1 ×10^6^ cells per mL in T cell media with IL-2. Unless otherwise indicated, CAR T cells were used for *in vitro* assays or transferred into mice on day 10-11 post activation.

### Retroviral transduction

Non-tissue culture treated 12-well plates were coated overnight at 4C with 1mL Retronectin (Takara) at 25ug/mL in PBS. Plates were washed with PBS and blocked with 2% BSA for 15min. Thawed retroviral supernatant was added at ∼1mL per well and centrifuged for 2 hours at 32C at 3200 RPM before the addition of cells.

### Cell Lines

The Kelly neuroblastoma, EW8 Ewing’s sarcoma, 143b and TC32 osteosarcoma cell lines were originally obtained from ATCC. In some cases, cell lines were stably transduced with GFP and firefly luciferase (GL). The CD19^+^CD22^+^ Nalm6-GL B-ALL cell line was provided by David Barrett. Nalm6-GD2 was created by co-transducing Nalm6-GL with cDNAs for GD2 synthase and GD3 synthase. A single cell clone was then chosen for high GD2 expression. Nalm6-22^KO^ and 22^low^ have been previously described and were kindly provided by Terry Fry. All cell lines were cultured in complete media (RPMI supplemented with 10% FBS, 10mM HEPES, 2mM GlutaMAX, 100 U/mL penicillin, and 100ug/mL streptomycin (Gibco)).

### Flow Cytometry

The anti-CD19 CAR idiotype antibody was kindly provided by Bipulendu Jena and Laurence Cooper^47^. The 1A7 anti-14G2a idiotype antibody was obtained from NCI-Frederick. CD22 and Her2 CARs were detected using human CD22-Fc and Her2-Fc recombinant proteins (R&D). The idiotype antibodies and Fc-fusion proteins were conjugated in house with Dylight488 and/or 650 antibody labeling kits (Thermo Fisher). T cell surface phenotype was assessed using the following antibodies:

From BioLegend: CD4-APC-Cy7 (clone OKT4), CD8-PerCp-Cy5.5 (clone SK1), TIM3-BV510 (clone F38-2E2), CD39-FITC or APC-Cy7 (clone A1), CD95-PE (clone DX2), CD3-PacBlue (clone HIT3a), From eBioscience: PD1-PE-Cy7 (clone eBio J105), LAG3-PE (clone 3DS223H), CD45RO-PE-Cy7 (clone UCHL1), CD45-PerCp-Cy5.5 (clone HI30), From BD: CD45RA-FITC or BV711 (clone HI100), CCR7-BV421 (clone 150503), CD122-BV510 (clone Mik-β3), CD62L-BV605 (clone DREG-56), CD4-BUV395 (clone SK3), CD8-BUV805 (clone SK1).

### Cytokine production

1×10^5^ CAR+ T cells and 1×10^5^ tumor cells were cultured in 200uL CM in 96-well flat bottom plates for 24 hours. For idiotype stimulation, serial dilutions of 1A7 were crosslinked in 1X Coating Buffer (BioLegend) overnight at 4C on Nunc Maxisorp 96-well ELISA plates (Thermo Scientific). Wells were washed once with PBS and 1×10^5^ CAR+ T cells were plated in 200uL CM and cultured for 24h. Triplicate wells were plated for each condition. Culture supernatants were collected and analyzed for IFNg and IL-2 by ELISA (BioLegend).

### Incucyte Lysis Assay

5×10^4^ GFP+ leukemia or 2.5×10^4^ GFP+ adherent tumor cells were co-cultured with CAR T cells in 200uL CM in 96-well flat bottom plates for up to 96 hours. Triplicate wells were plated for each condition. Plates were imaged every 2-3 hours using the IncuCyte ZOOM Live-Cell analysis system (Essen Bioscience). 4 images per well at 10X zoom were collected at each time point. Total integrated GFP intensity per well was assessed as a quantitative measure of live, GFP+ tumor cells. Values were normalized to the starting measurement and plotted over time. E:T ratios are indicated in the Figure legends.

### Western Blotting and Immunoprecipitations

Whole-cell protein lysates were obtained in nondenaturing buffer (150 mmol/L NaCl, 50 mmol/L Tris-pH8, 1% NP-10, 0.25% sodium deoxycholate). Protein concentrations were estimated by Bio-Rad colorimetric assay. Immunoblotting was performed by loading 20μg of protein onto 11% PAGE gels followed by transfer to PVF membranes. Signals were detected by enhanced chemiluminescence (Pierce) or with the Odyssey imaging system. Representative blots are shown. The following primary antibodies used were purchased from Cell Signaling: c-Jun (60A8), P-c-Jun^Ser73^ (D47G9), JunB(C37F9), BATF(D7C5) and IRF4(4964). The BATF3 (AF7437) antibody was from R&D. Immunoprecipitations were performed in 100mg of whole-cell protein lysates in 150μL of nondenaturing buffer and 7.5mg of agar-conjugated antibodies c-Jun (G4) or JunB (C11) (Sant Cruz Biotechnology). After overnight incubation at 4°C. Beads were washed 3 times with nondenaturing buffer, and proteins were eluted in Laemmli sample buffer, boiled, and loaded onto PAGE gels. Detection of immunoprecipitated proteins was performed with above-mentioned reagents and antibodies.

### Mice

Immunocompromised NOD/SCID/IL2Rγ^−/−^ (NSG) mice were purchased from JAX and bred in-house. All mice were bred, housed, and treated under Stanford University IACUC (APLAC) approved protocols. 6-8 week old mice were inoculated with either 1×10^6^ Nalm6-GL leukemia via intravenous (IV) or 0.5-1×10^6^ 143B osteosarcoma via intramuscular (IM) injections. All CAR T cells were injected IV. Time and treatment dose are indicated in the Figure legends. Leukemia progression was measured by bioluminescent imaging using the IVIS imaging system. Values were analyzed using Living Image software. Solid tumor progression was followed using caliper measurements of the injected leg area. 5 mice per group were treated in each experiment, and each experiment was repeated 2 or 3 times as indicated. Mice were randomized to ensure equal pre-treatment tumor burden before CAR T cell treatment.

### Blood and tissue analysis

Peripheral blood sampling was conducted via retro-orbital blood collection under isoflurane anesthesia at the indicated time points. 50µL blood was labeled with CD45, CD3, CD4, and CD8, lysed using BD FACS Lysing Solution and quantified using CountBright Absolute Counting beads (Thermo Fisher) on a BD Fortessa flow cytometer.

### ATAC-seq

ATAC-seq library preparation was carried out as described previously^48^. Briefly, 100,000 cells from each sample were sorted by FACS into CM, centrifuged at 500g at 4°C, then resuspended in ATAC-seq Resuspension Buffer (RSB) (10 mM Tris-HCl, 10 mM NaCl, 3mM MgCl2) supplemented with 0.1% NP-40,0.1% Tween-20, and 0.01% Digitonin. Samples were split into two replicates each prior to all subsequent steps. Samples were incubated on ice for 3 minutes, then washed out with 1 mL RSB supplemented with 0.1% Tween-20. Nuclei were pelleted at 500g for 10 minutes at 4°C. The nuclei pellet was resuspended in 50 μL transposition mix (25 μl 2x TD buffer, 2.5 μl transposase (Illumina), 16.5 μl PBS, 0.5 μl 1% digitonin, 0.5 μl 10% Tween-20, 5 μl H2O) and incubated at 37°C for 30 minutes in a thermomixer with 1000 RPM shaking. The reaction was cleaned up using the Qiagen MinElute PCR Purification Kit. Libraries were PCR-amplified using the NEBNext Hi-Fidelity PCR Master Mix and custom primers (IDT) as described previously^20^. Libraries were sufficiently amplified following 5 cycles of PCR, as indicated by qPCR fluorescence curves^20^. Libraries were purified with the Qiagen MinElute PCR Purification Kit and quantified with the KAPA Library Quantification Kit. Libraries were sequenced on the Illumina NextSeq at the Stanford Functional Genomics Facility with paired-end 75bp reads. Adapter sequences were trimmed using SeqPurge and aligned to hg19 genome using bowtie2. These reads were then filtered for mitochondrial reads, low mapping quality (Q >=20), and PCR duplicates using Picard tools. Then we converted the bam to a bed and got the Tn5 corrected insertion sites (“+” stranded + 4 bp, “-” stranded −5 bp). To Identify peaks, we called peaks for each sample using MACS2 “--shift −75 --extsize 150 --nomodel --call-summits --nolambda -- keep-dup all -p 0.00001” using the insertion beds. To get a union peak set, we (1) extended all summits to 500bp, (2) merged all summit bed files and then (3) used bedtools cluster and selected the summit with the highest MACS2 score. This was then filtered by the ENCODE hg19 blacklist (https://www.encodeproject.org/annotations/ENCSR636HFF/), to remove peaks that extend beyond the ends of chromosomes. We then annotated these peaks using HOMER and computed the occurrence of a TF motif using motifmatchr in R with chromVARMotifs HOMER set. To create sequencing tracks, we read the Tn5 corrected insertion sites into R and created a coverage pileup binned every 100bp using rtracklayer. We then counted all insertions that fell within each peak to get a counts matrix (peak x samples). To determined differential peaks we first used peaks that were annotated as “TSS” as control genes or “Housekeeping Peaks” for DESeq2 and then computed differential peaks with this normalization. All clustering was performed using the regularized log transform values from DESeq2. Transcription factor motif deviation analysis was carried out using chromVAR as described previously^21^. TF motif enrichment were calculated using a hypergeometric test in R testing the representation of a motif (from motifmatchr above) in a subset of peaks vs all peaks.

### Subset RNA-seq

For T cell subset-specific RNA-seq, T cells were isolated from healthy donor buffy coats as described above. Before activation, naïve and central memory CD4+ or CD8+ subsets were isolated using a BD FACSAria cell sorter (Stem Cell FACS Core, Stanford University School of Medicine) using the following markers: Naïve (CD45RA+CD45RO-, CD62L+, CCR7+, CD95-, and CD122-), Central Memory (CD45RA-CD45RO+, CD62L+, CCR7+). Sorted starting populations were activated, transduced, and cultured as described above. On days 7, 10, and 14 of culture, CAR+ CD4+ and CD8+ cells were sorted, and RNA was isolated using Qiagen mRNEasy kit. Samples were library prepped and sequenced via Illumina NextSeq paired end platform by the Stanford Functional Genomics Core.

### Bulk RNA-seq

For bulk RNA isolation, healthy donor T cells were prepared as described. On day 10 or 11 of culture, total mRNA was isolated from 2 x 10^6^ bulk CAR T cells using Qiagen RNEasy Plus mini isolation kit. Bulk RNA-seq was performed by BGI America (Cambridge, MA) using the BGISEQ-500 platform, single end 50bp-read length, at 30 x 10^6^ reads per sample. Principal component analysis was performed using stats package and plots with ggplot2 package in R (version 3.5)^49^. Gene set enrichment analysis was performed using the GSEA software (Broad Institute) as described^50, 51^.

### Single Cell RNA-seq

To compare gene expression in single CD19-CAR and GD2-CAR T cells, we sorted naïve T-cell subset on day 0 for subsequent single-cell analysis on day 10 using the Chromium platform (10x Genomics) and the Chromium Single Cell 3’ v2 Reagent Kit according to the manufacturer’s instructions. cDNA libraries were prepared separately for CD19-CAR and GD2-CAR cells, and the CD4+ cells and CD8+ cells were combined in each run to be separated bioinformatically downstream. Sequencing was performed on the Illumina NextSeq system (paired-end, 26 bp into read 1 and 98 bp into read 2) to a depth >100,000 reads per cell. Single-cell RNA-sequencing reads were aligned to the Genome Reference Consortium Human Build 38 (GRCh38), normalized for batch effects, and filtered for cell events using the *Cell Ranger* software (10X Genomics). A total of 804 CD19-CAR and 726 GD2-CAR T cells were sequenced to an average of 350,587 post-normalization reads per cell. The cell-gene matrix was further processed using the *Cell Ranger R Kit* software (10X Genomics) as described^52^. Briefly, we first selected genes with ≥1 unique molecular identifier (UMI) counts in any given cell. UMI counts were then normalized to UMI sums for each cell and multiplied by a median UMI count across cells. Next, the data were transformed by taking a natural logarithm of the resulting data matrix.

### Statistical Analysis

Unless otherwise noted, statistical analyses for significant differences between groups were conducted using upaired 2-tailed t-tests without assuming consistent SD using GraphPad Prism 7. For bulk RNA-seq in Figure 2C, the nonparametric Wilcoxon matched-pair signed rank test was used. Survival curves were compared using the Log-rank Mantel-Cox test. A table with the full statistical analysis, including exact p values, t ratio, and dof can be found in the supplementary materials.

### Data Availability

The sequencing datasets generated during the current study will be made publicly available upon acceptance and prior to final publication.

## ACKNOWLEDGEMENTS

This work was supported by a Stand Up To Cancer–St. Baldrick’s–National Cancer Institute Pediatric Dream Team Translational Cancer Research Grant (C.L.M.). This work was supported by Parker Institute for Cancer Immunotherapy (C.L.M., H.Y.C.), and NIH P50-HG007735 (H.Y.C.). H.Y.C. is an Investigator of the Howard Hughes Medical Institute. A.T.S. was supported by a Parker Bridge Scholar Award from the Parker Institute for Cancer Immunotherapy and a Career Award for Medical Scientists from the Burroughs Welcome Fund. R.C.L. was supported by the Emerson Collective Cancer Research Fund.

## AUTHOR CONTRIBUTIONS

R.C.L. cloned the constructs, designed and performed experiments, analyzed data, and wrote the manuscript. E.W.W. and E.S. designed and performed experiments. D.G., J.G., A.T.S., and H.Y.C. performed and analyzed ATAC-seq. Z.G., C.D.B., and S.R.Q. performed/analyzed single cell RNA-seq. H.A. and R.J. performed/analyzed bulk RNA-seq. V.T. cloned the JUN-DD constructs and performed experiments. R.M. cloned the HA-GD2 CAR. P.X. did animal injections and imaging. C.L.M. designed experiments and wrote the manuscript.

## EXTENDED DATA FIGURES

**Extended Data Figure 1:**
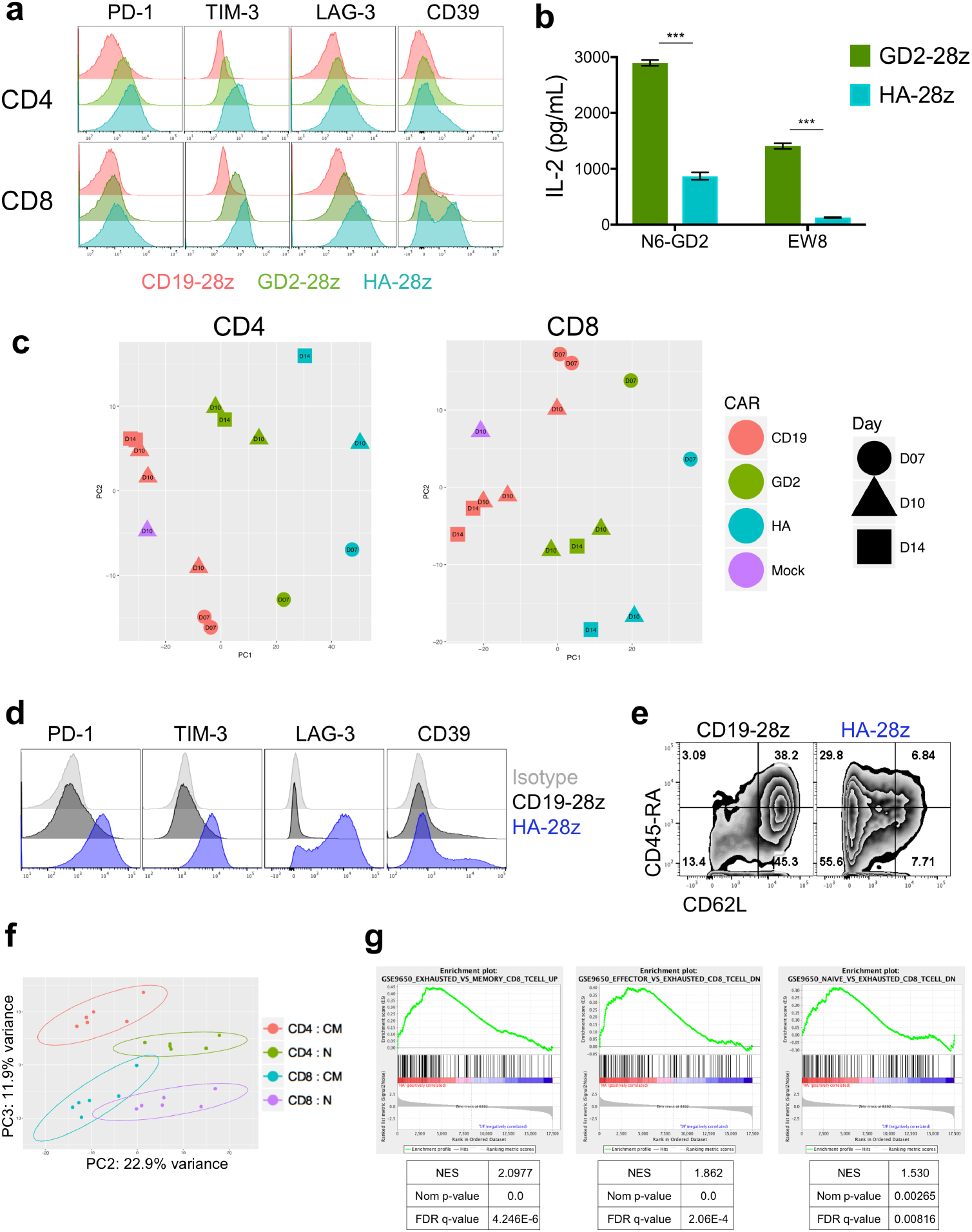
High Affinity (HA) 14g2a-GD2^E101K^ CAR T cells manifest an exaggerated exhaustion signature compared to the original 14g2a-GD2 CAR. **a)** Surface inhibitory receptor expression in CD19, GD2, and HA-GD2^E101K^ CAR T cells at day 10 of culture. High affinity E101K mutation results in increased inhibitory receptor expression in CD4^+^ and CD8^+^ CAR T cells, compared to parental GD2 CAR. **b)** IL-2 secretion following 24h co-culture of HA-GD2^E101K^ or original GD2-28z CAR T cells with GD2^+^ target cells. The increased exhaustion profile of HA-GD2^E101K^ CAR T cells corresponds to decreased functional activity, as measured by the ability to produce IL-2 upon stimulation. Error bars represent mean ± SD of triplicate wells. Representative of at least 4 independent experiments with similar results. **c)** PCA of bulk RNA-seq demonstrates larger variance between HA-GD2^E101K^ and CD19 CAR T cells, whereas GD2-28z(sh) CAR T cells are intermediary. Left – CD4^+^ T cells. Right – CD8^+^ CAR T cells, Naïve-derived. d-e) HA-GD2^E101K^ CAR expression causes enhanced inhibitory receptor expression **(d)** and decreased memory formation **(e)** in CD4^+^ CAR T cells. (CD8^+^ data in Figure 1). **f)** RNA-seq PCA from Figure 1e showing PC2 separation is driven by CM vs N and PC3 separation driven by CD4 vs CD8. **g)** GSEA: gene sets upregulated in day 10 HA-28z CAR T cells vs CD19-28z CAR T cells showed significant overlap with genes upregulated in Exhausted vs Memory CD8^+^ (left), Exhausted vs Effector CD8^+^ (middle), and Exhausted vs Naïve CD8^+^ (right) in a mouse model of chronic viral infection (Wherry et al. Immunity, 2007). * p < .05, ** p < .01, *** p < .001. PCA – principle component analysis, NES – normalized enrichment score.

**Extended Data Figure 2:**
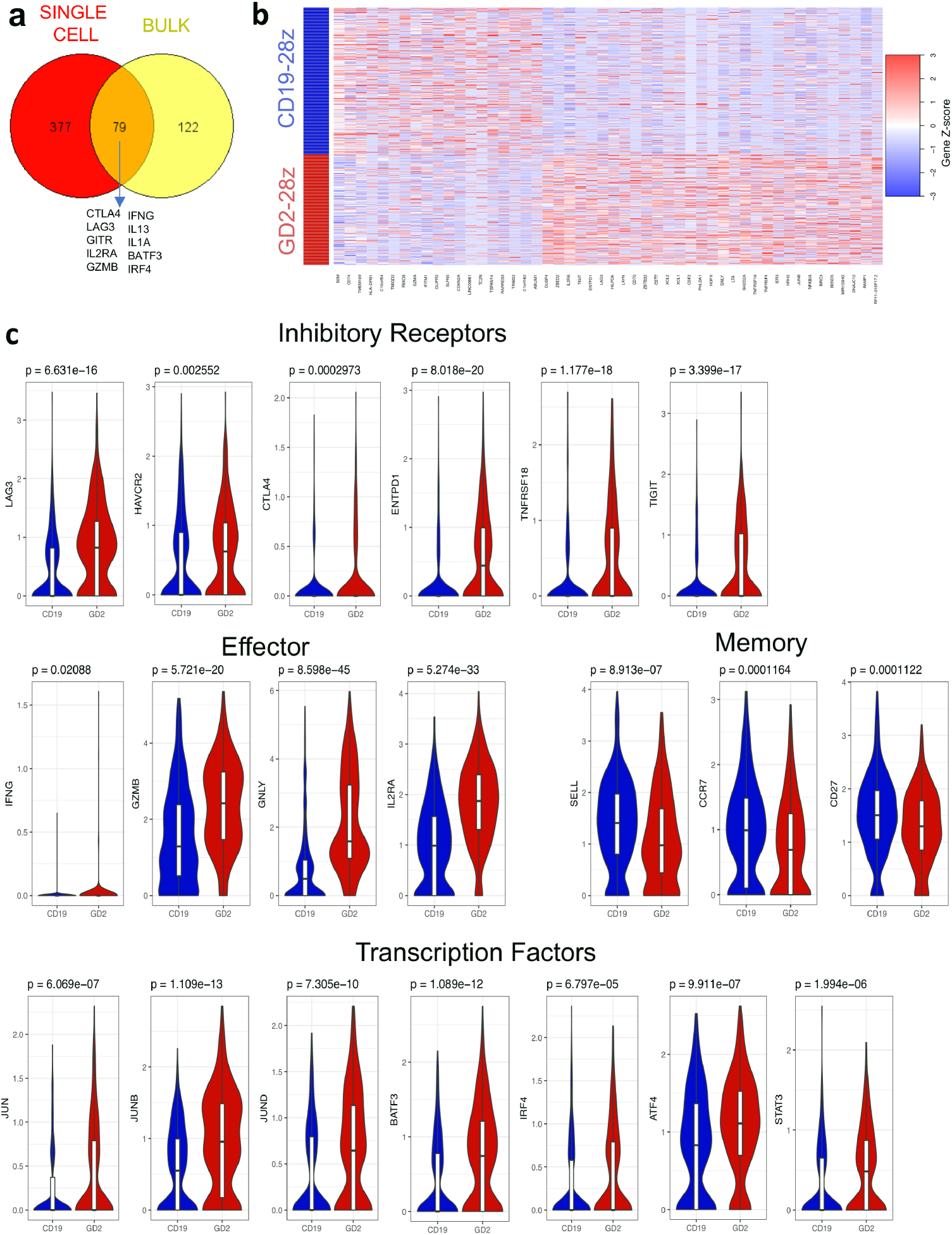
GD2-28z CAR T cells display an exhaustion signature at the single cell level. **a)** Venn diagram showing overlapping genes in differential expression analysis of single cell data (red) and the top 200 genes driving the separation of CD19 and HA-28z CAR T cells in bulk RNA-seq (yellow, Figure 1f). 79 out of the top 200 genes from bulk RNA-seq are differentially expressed by DESeq2 analysis in GD2-28z vs CD19-28z single cells. Highlighted genes from the intersection include inhibitory receptors (*CTLA4*, *LAG3*, *GITR*, effector molecules *CD25*, *IFNG*, *GZMB*, and cytokines *IL13* and *IL1A* and AP-1/bZIP family transcription factors *BATF3* and *IRF4*. **b)** Heatmap clustering the top 50 differentially expressed genes in GD2-28z vs CD19-28z single cell transcriptome analysis. Each row represents one cell. **c)** Violin plots depicting individual gene expression in CD8^+^ GD2-28z and CD19-28z single CAR T cells. Genes upregulated in GD2 CAR T cells include inhibitory receptors, effector molecules, and AP-1 family transcription factors, while CD19 CAR T cells have increased expression of memory-associated genes. *P*-values that are displayed for each gene above the individual plots were calculated using unpaired two-tailed Wilcoxon-Mann-Whitney *U* test.

**Extended Data Figure 3:**
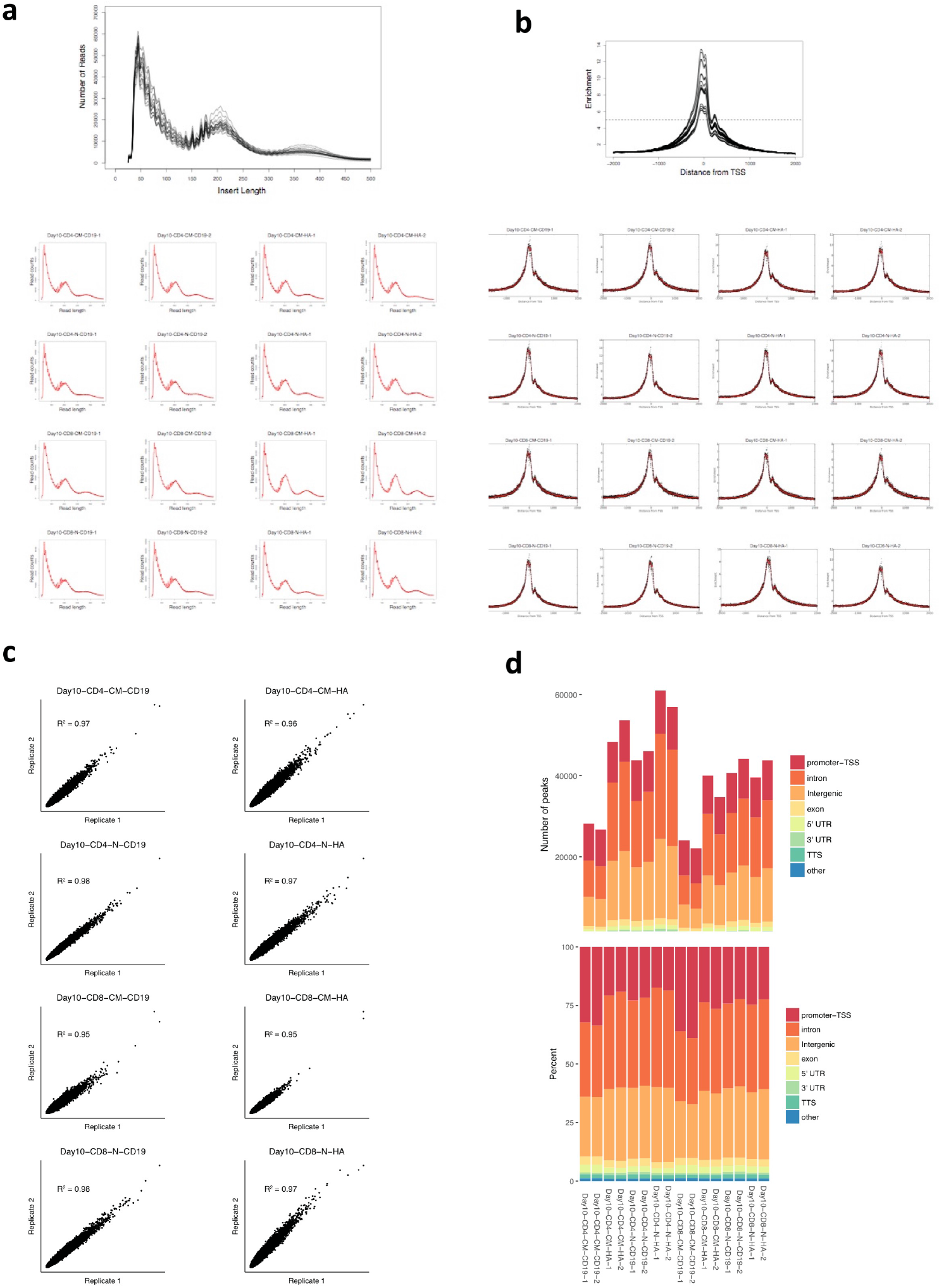
ATAC-seq data quality control. **a)** Insert length **b)** insert distance from transcriptional start site (TSS) for combined (top) and individual samples (below). **c)** Correlation between replicate samples. **d)** Location of mapped peaks in each sample by total number of peaks (upper) and frequency of total (lower).

**Extended Data Figure 4:**
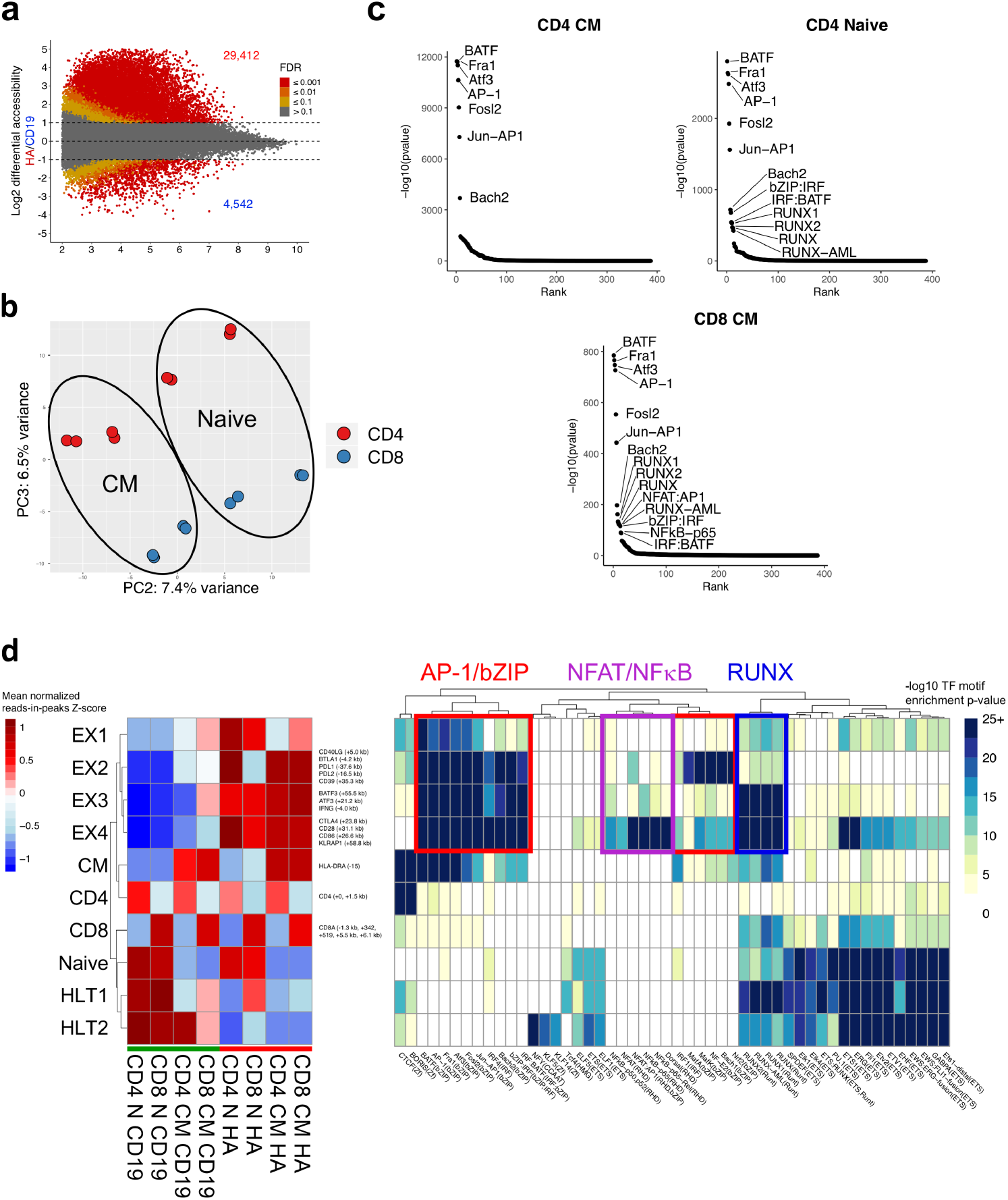
AP-1 family comprise the most significantly enriched transcription factor motifs in HA-28z exhausted CAR T cells. **a)** Differentially accessible chromatin regions in CD4^+^ CD19 and HA CAR T cells (D10). Both N and CM subsets are incorporated for each CAR. **b)** PCA from Figure 1h showing PC2 separation is driven by CM vs N and PC3 separation driven by CD4 vs CD8. **c)** Top transcription factor motifs enriched in chromatin regions differentially accessible in HA-28z CAR T cells comprise AP-1/bZIP family factors in all starting T cell subsets. CD8^+^ Naïve subset is shown in Figure 2. **d)** Peak clustering by shared regulatory motif (left) and enrichment heat map of transcription factor motifs (right) in each cluster. 10 different clusters including clusters associated with exhausted (EX1-EX4) or healthy (HLT1-HLT2) CAR T cells, CM (CM) or N (Naive) starting subset, and CD4 or CD8 T cell subset. Genes of interest in each cluster are highlighted to the right. (N – naïve, CM – central memory).

**Extended Data Figure 5:**
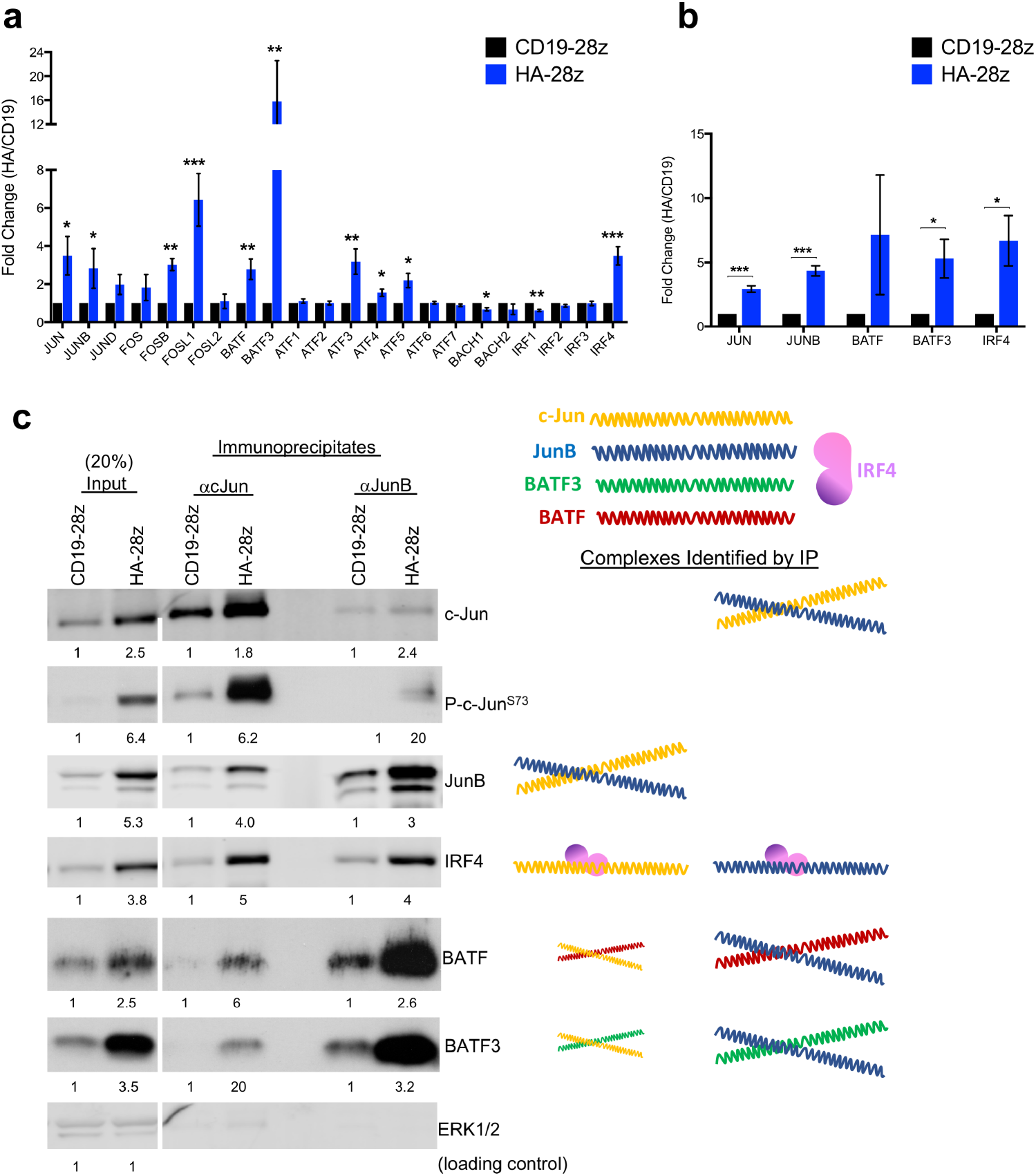
AP-1/bZIP family transcription factors are upregulated in HA-28z CAR T cells and form immunoregulatory complexes. **a)** Fold change in the gene expression (HA/CD19) for the indicated AP-1/bZIP and IRF family genes from RNA sequencing data from Figure 2. Error bars represent mean ± SEM of n = 6 samples across 3 independent donors. **b)** Fold change in the protein expression (HA/CD19) for the indicated AP-1/bZIP and IRF family proteins was determined by densitometry analysis of western blots. Error bars represent mean ± SEM of n = 4 experiments across 3 independent donors. * p < .05, ** p < .01, *** p < .001. **c)** Western blot analysis for the indicated AP-1/bZIP and IRF family member proteins at input (left columns) or after immunoprecipitation for c-Jun (middle columns) or JunB (right columns) in CD19 and HA-28z CAR T cells. Numbers below represent the fold increase in protein expression for HA vs CD19 at each condition and colored shapes represent the complexes identified sized to scale. IP-western blots demonstrate the increased presence of c-Jun/JunB, c-Jun/IRF4, c-Jun/BATF, and c-Jun/BATF3 complexes in HA-28z CAR T cells. IRF4 is also bound at a similar ratio to JunB, while BATF and BATF3 show a preferential complexing with JunB.

**Extended Data Figure 6:**
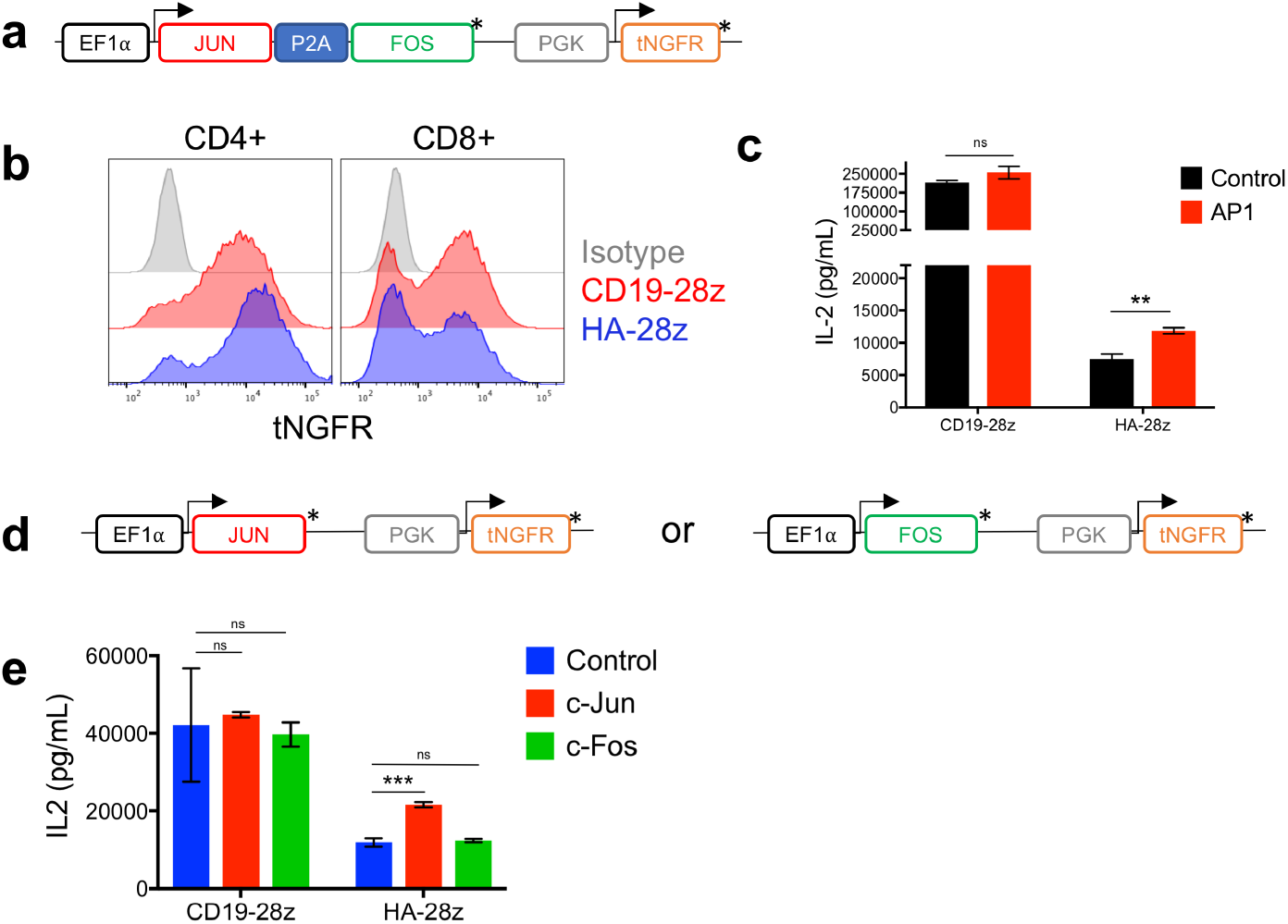
Enhanced activity of AP1-modified CAR T cells is dependent on c-Jun but not c-Fos. **a-c)** CAR T cells were co-transduced with (AP1) or without (Control) a lentiviral vector encoding both AP1 transcription factors Fos and Jun and a truncated NGFR (tNGFR) surface selection marker. **a)** Schematic of the lentiviral construct. **b)** Representative transduction efficiency of AP1 modified CAR T cells as measured by NGFR surface expression in indicated CD4^+^ and CD8^+^ CAR T cells. **c)** IL-2 production in control (black) or AP1-modified (red) CAR T cells following 24hr stimulation with 143B-19 target cells. AP1-modified HA-28z CAR T cells show increased IL-2 production compared to control CAR T cells. Representative experiment of 2 independent experiments with similar results. **d-e)** CAR T cells were co-transduced with lentiviral vectors encoding either AP1 transcription factor Fos or Jun and a truncated NGFR (tNGFR) surface selection marker. **d)** Schematics of the Fos and Jun lentiviral constructs. **e)** IL-2 production in control (blue), Fos (green), or Jun (red) modified CAR T cells following 24hr stimulation with Nalm6-GD2 target cells. Error bars represent mean ± SD of triplicate wells. Representative experiment of 2 independent experiments with similar results. In a and d, * denotes a stop codon. * p < .05, ** p < .01, *** p < .001, ns p > 0.05.

**Extended Data Figure 7:**
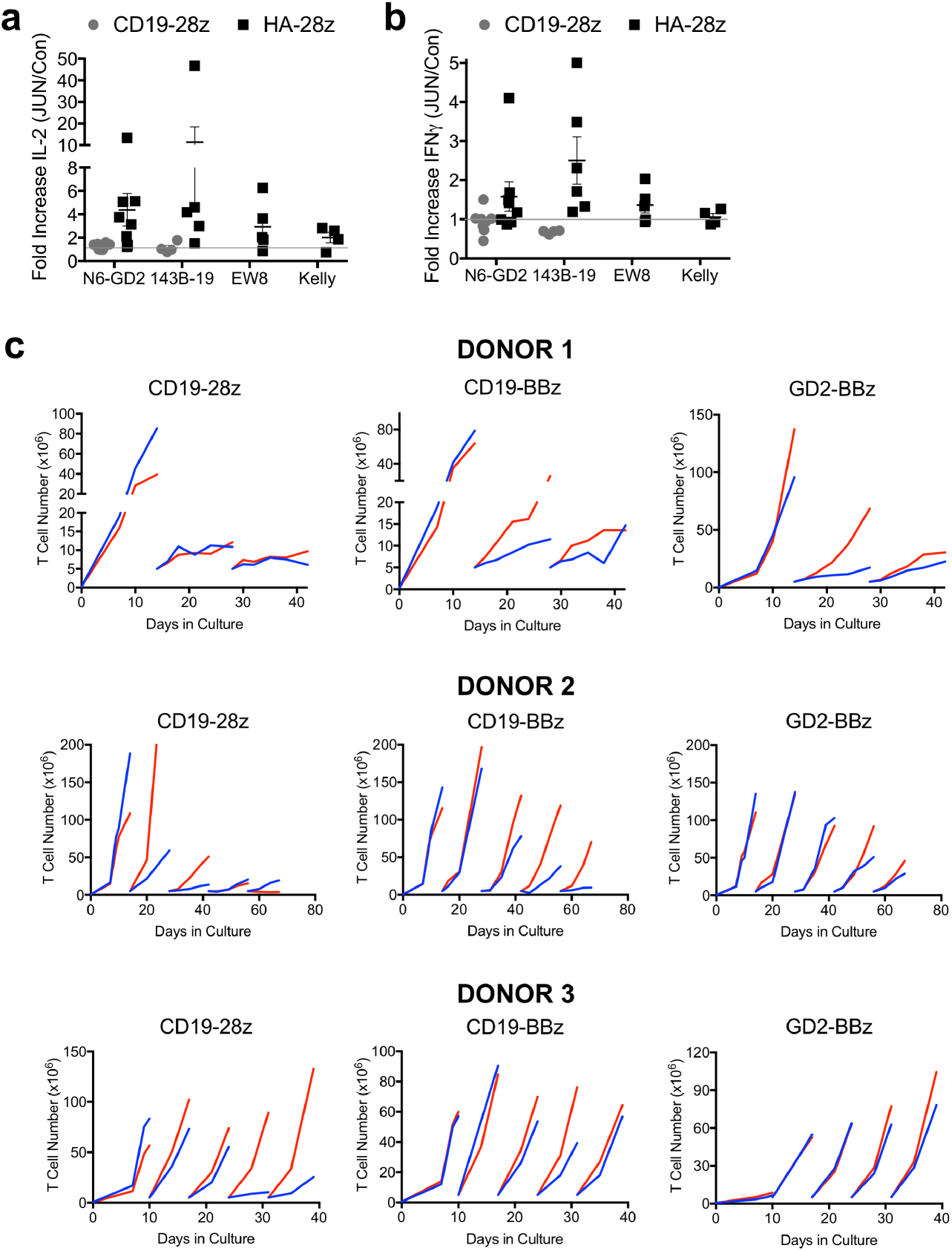
Extended functional assessment of JUN-modified CAR T cells. **a-b)** Fold increase in IL-2 **(a)** and IFNγ **(b)** release following 24hr co-culture with the indicated target cells in JUN vs Control CD19 and HA-28z CAR T cells. Each dot represents one independent experiment from different donors. **c)** Extended expansion of control (blue) or JUN-modified (red) CAR T cells in vitro in 3 independent experiments with 3 different healthy donors. At the indicated time points, T cells were re-plated in fresh T cell media + 100 IU/mL IL2. T cells were counted and fed to keep cells at 0.5×10^6^/mL every 2-3 days. For DONOR-1, 5×10^6^ viable T cells were re-plated on days 14 and 28. For DONOR-2, 5×10^6^ viable T cells were re-plated on days 14, 28, 42, and 56. For DONOR-3, 5×10^6^ viable T cells were re-plated on days 10, 17, 24, and 31.

**Extended Data Figure 8:**
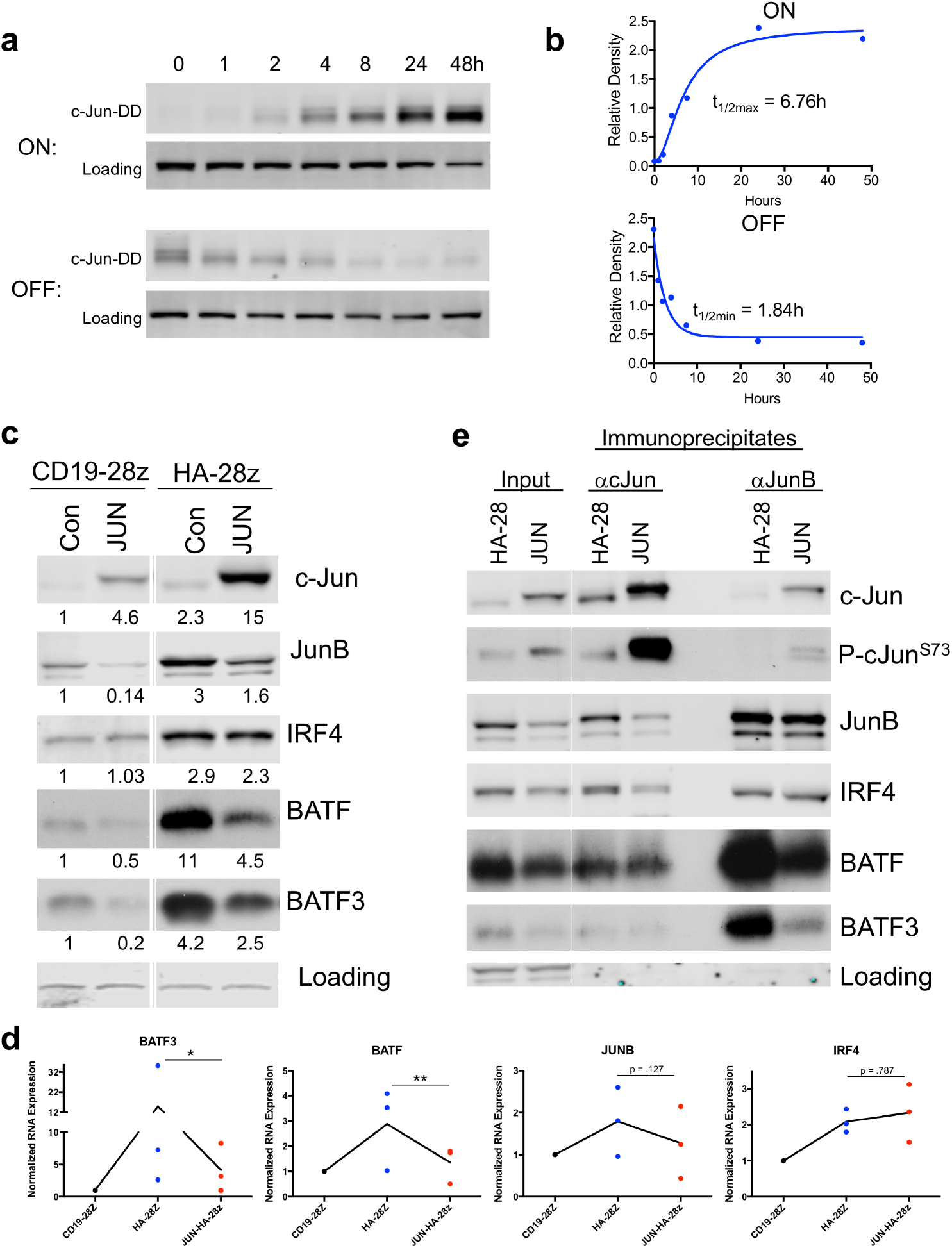
c-Jun overexpression decreases the prevalence and complexing of inhibitory AP-1 family members JunB, BATF, and BATF3. **a)** Kinetics of drug-induced c-Jun stability in JUN-DD CAR T cells as assessed by western blot. At time 0, 10uM TMP was either added to untreated cells (ON) or washed out of previously treated cells (OFF). Cells were removed from each condition at 1, 2, 4, 8, 24, and 48hr and prepared for western blot analysis of c-Jun expression. The observed band corresponds to the size of JUN-DD. **b)** Densitometry analysis was performed on the blots from (a) and normalized to a loading control. Expression was plotted vs time and first order kinetics curves fit to the data to determine t1/2 for OFF and ON kinetics. **c)** Western blot analysis for the indicated AP-1/bZIP and IRF family member proteins in control and JUN-CAR T cells (D10). Numbers below represent the fold change in protein expression compared to CD19. **d)** Corresponding decrease in mRNA expression of *BATF*, *BATF*3, and *JUNB* in JUN-HA-28z CAR T cells compared to HA-28z. n=3 donors, normalized to CD19 mRNA. **e)** c-Jun overexpression decreases inhibitory JunB/BATF and JunB/BATF3 complexes by IP-western blot analysis. Input (left columns), immunoprecipitation for c-Jun (middle columns), or JunB (right columns) in Control or JUN-HA-28z CAR T cells. IRF4 protein, mRNA, and complexing with c-Jun is unchanged.

**Extended Data Figure 9:**
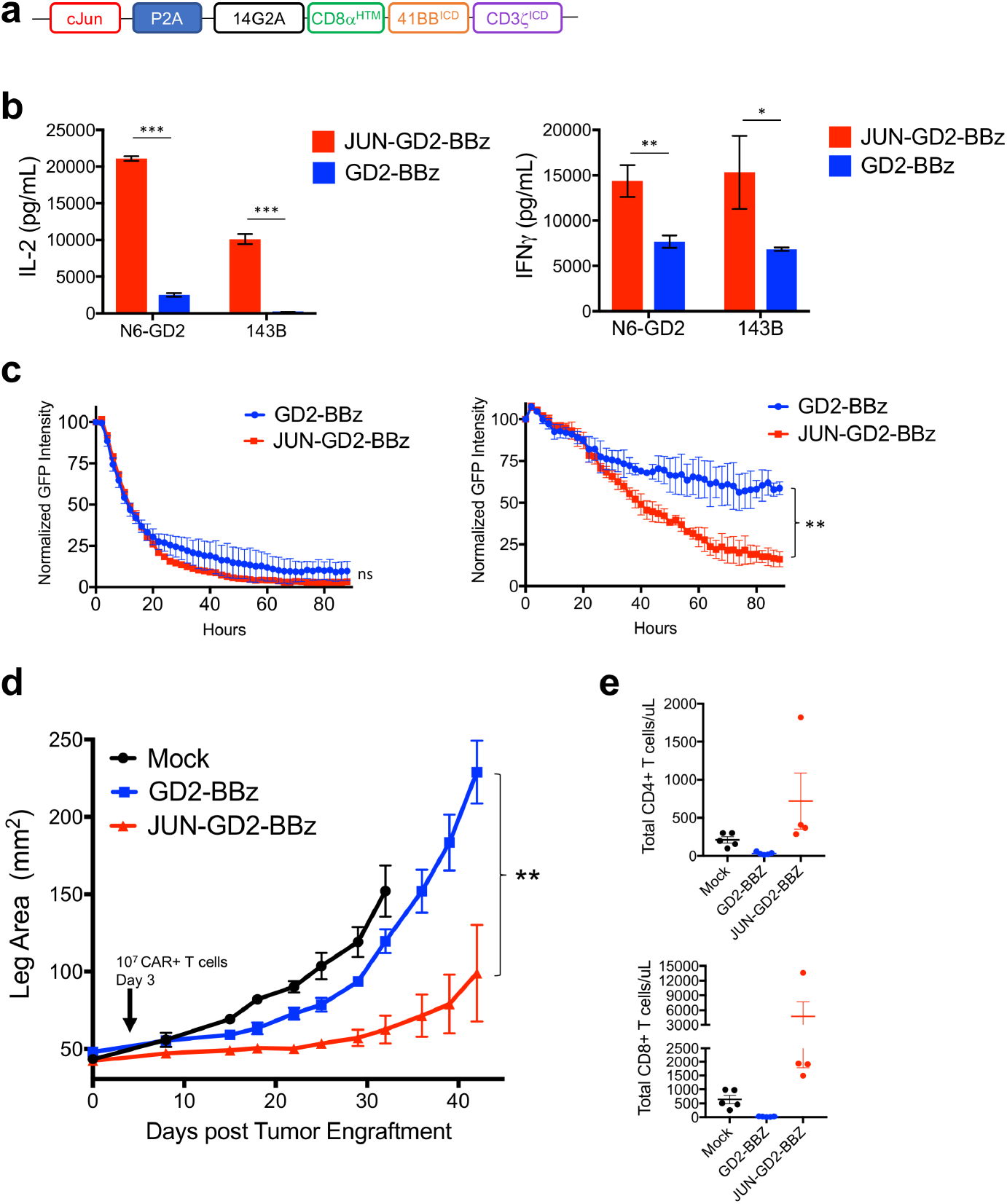
JUN-CAR T cells enhance GD2-BBz CAR T cell function in solid tumors. **a)** Vector schematic of JUN-GD2-BBz retroviral vector construct. **b)** IL-2 (left) and IFNγ (right) production in JUN-modified (red) or control (blue) GD2-BBz CAR T cells following 24hr stimulation with Nalm6-GD2 or 143B target cells. **c)** GD2-BBz CAR T cell lysis of GFP+ Nalm6-GD2 target cells at 1:1 (left) or 1:4 (right) E:T ratio. In a-c, error bars represent mean ± SD of triplicate wells. Representative of at least 3 independent experiments. In **d-e**, NSG mice were inoculated with 0.5×10^6^ 143B-19 osteosarcoma cells via intramuscular injection. 1×10^7^ Mock, GD2-BBz, or JUN-GD2-BBz CAR T cells were given IV on day 3. **d)** Tumor growth was monitored by caliper measurements. **e)** Peripheral blood CD4^+^ (upper) or CD8^+^ (lower) T cell counts at day 14 post tumor engraftment. Error bars represent mean ± SEM of n=5 mice per group. Representative of 2 independent experiments although early deaths (unrelated to tumor size) precluded survival curves in both models. * p < .05, ** p < .01, *** p < .001. HTM – hinge/transmembrane. ICD – intracellular domain.

